# Dietary fibre combinations modulate gut microbiome composition and metabolic responses in zebrafish, with lasting effects on intestinal barrier gene expression

**DOI:** 10.64898/2026.11.06.764917

**Authors:** Justin Abi Assaf, Gwenaelle Le Gall, Ruth Sullivan, Anastasia Bezrucico-Pyatakov, Daniel Clifton, Isaac Harris, Özlem Pullin, Simone Immler

## Abstract

Food sources provide a complex mixture of dietary fibres, not a single type. Consuming a diverse range of these dietary fibre-containing foods is associated with health benefits. However, how the effects of combining dietary fibres translate across different biological levels remains poorly characterised. This study uses an integrated microbiome-metabolome analytical approach that combines 16S rRNA gene sequencing with metabolomic profiling of the intestine, brain and liver. Specifically, we examined the combined effects of apple pectin (AP) and cellulose (C) within the same diet, compared with their isolated effects, in adult female and male zebrafish (*Danio rerio*). Notably, the combined apple pectin and cellulose diet (APC) produced a microbiome composition that could not be predicted by simply adding the changes observed with each dietary fibre individually, indicating a significant non-additive interaction between the two dietary components, specifically a less-than-additive response in gut microbiome composition. The resulting community composition under the combined diet also more closely resembled that induced by apple pectin. We found that predicted microbial functional capabilities did not translate directly into quantified intestinal and brain short-chain fatty acid (SCFA) concentrations, nor did intestinal SCFA levels directly mirror those observed in the brain. Moreover, we used targeted qPCR gene expression analysis to assess responses to a subsequent saponin challenge after dietary fibre cessation, revealing a potentially protective effect on the intestinal barrier. Collectively, we show that dietary fibres within the same diet interact to shape the gut microbiome and do not act in isolation, leading to non-linear health ramifications for the host that may persist even after active consumption ends.

**Graphical Abstract:** 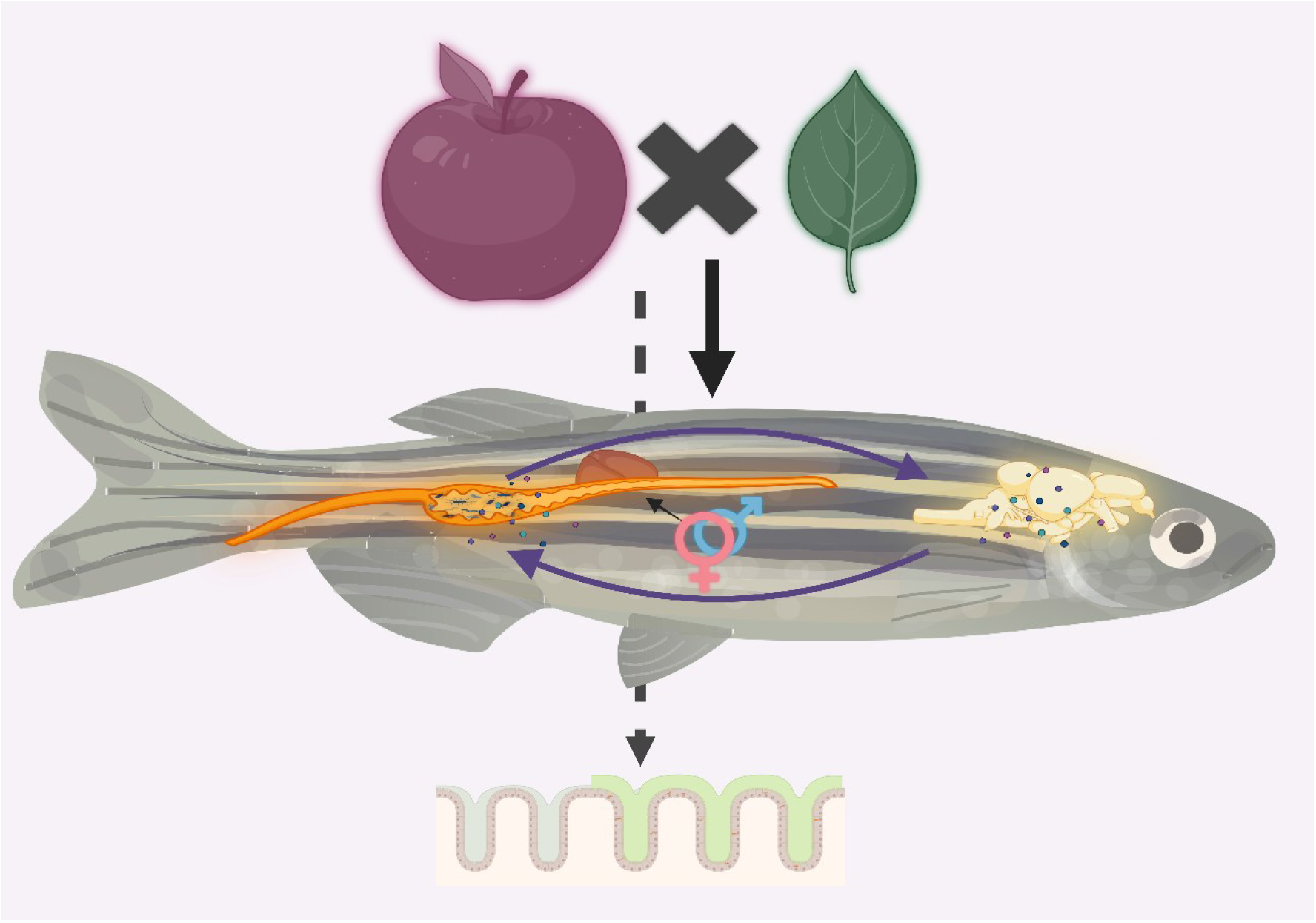

Created in https://BioRender.com

## INTRODUCTION

Dietary fibres (DFs) are seldom consumed in isolation, but rather as part of complex diets containing multiple structurally and functionally heterogeneous DF sources. The biological effects of DFs depend not only on the properties of individual components, but also on the host-microbiome context in which they are consumed^1,2^. Moreover, because their physicochemical properties drive interactions with gut microbes, DF effects can extend beyond their own microbial breakdown. This fosters interactions among other dietary components and substrates, as well as among members of the gut microbial community^2^, leading to potential downstream health ramifications. We have previously shown^3^ that DF effects extend beyond physicochemical properties and depend on the broader dietary context, including dosage and feeding frequency. However, intra-dietary fibre interactions remain underexplored, particularly the effects of mixed DF exposure, which better reflects real-life dietary patterns^4,5^. Importantly, these dietary interactions also occur within a host-dependent biological context. Our earlier work^3^ showed that DF effects also depend on the host’s intrinsic and biological conditions. Thus, the host plays a key role in shaping both the baseline composition of the gut microbiome and the metabolic outputs derived from the breakdown of different DFs^2^. Consequently, the gut microbiome sits at the interface between host biology and dietary exposure^6^. It continually responds to and adapts to changes in both dietary exposure and the host environment. This capacity to respond and adapt to perturbation is captured by the concept of ecological resilience, which refers to the gut microbiome’s capacity to withstand perturbations or to recover after disruption^7,8^. In our context, dietary changes induced by different DFs may shift the microbial community to a new state, followed by adaptation or a return to its baseline state once the perturbation or stimulus is removed. However, recovery is not always complete, and microbial communities may instead stabilise in an altered state following perturbation^7,9^. Importantly, recovery of the gut microbiome does not necessarily imply complete host restoration. Although the gut microbiome may return towards its pre-intervention state after a temporary dietary change ends, emerging evidence suggests that prior environmental and dietary exposures can leave persistent imprints on overall metabolism^10^ and the intestinal mucosa^11^. For example, intestinal epithelial and stem cells can retain epigenetic changes following previous dietary or inflammatory exposure, potentially modifying their response to subsequent stress or challenges^12^. Thus, prior DF intake may modulate the host-microbiome and leave a lasting immunological or structural imprint on intestinal barrier function, thereby altering the response to subsequent intestinal challenges.

In this study, we characterised putative interactions between different DFs within the same diet, compared with the effect of each DF in isolation. We designed and conducted a 2 × 2 factorial dietary intervention in which we fed four different diets to adult zebrafish reared in mixed-sex cohorts. The diets included the Control basal diet, an apple pectin diet (15AP), a cellulose diet (15C), and a diet containing equal amounts of apple pectin and cellulose combined (30APC) (**Fig. S1**). We fed these diets twice daily for three weeks (21 days). Then, using 16S rRNA sequencing, we examined how these diets shaped gut microbiome composition and diversity, as well as the predicted microbial metabolic pathways. Having previously established that short-chain fatty acids (SCFAs) can be quantified using targeted metabolomics (LC-MS/MS) in both the zebrafish intestine and brain^3^, we next asked how DF-induced changes in gut microbial composition translate into intestinal SCFA concentrations. We also explored whether intestinal levels mirror brain SCFA concentrations to understand SCFA dynamics along the host gut-brain axis. To provide a comprehensive overview of host metabolism, we also screened the liver for a broad range of metabolites, covering both primary metabolism and microbially derived metabolites. The initial dietary intervention experiment allowed us to investigate the host microbiome-metabolism crosstalk in response to DFs. However, we also wanted to test whether the host could retain some phenotypic improvements after DF cessation. Thus, in the second phase of the experiment (**Fig. S2**), we orally administered purified saponin, an established intestinal irritant in fish^13^, to challenge the host before testing gene expression related to both immune response and intestinal barrier function.

We show that apple pectin and cellulose, when combined, can elicit a gut microbiome response that differs from responses when administered in isolation and does not align with the predicted direction, indicating a significant non-additive interaction between the two DFs. Moreover, we found that shifts in microbial composition and predicted function do not always translate into quantifiable changes in tissue metabolite availability, highlighting the host’s role in fine-tuning how DF-microbiome interactions are ultimately reflected at the tissue level. In tandem, we show that the health ramifications of DF intake may extend beyond the period of dietary exposure, with effects persisting at the gut barrier and potentially attenuating the response to a subsequent stressor.

We already have clear evidence of the health benefits of DFs^14^. Yet, we still do not fully understand the mechanisms or the conceptual framework by which they elicit host responses that mitigate disease risk^2^. Our data highlight the complex, highly dynamic triadic crosstalk among the elements of the diet-microbe-host complex. We show that the overall health ramifications of DFs are not linear and cannot be readily inferred from a DF’s identity alone. Instead, DF-mediated effects arise from heterogeneity in their physicochemical properties, the presence of other dietary substrates, and the complexity of the gut microbiome and host physiological factors.

## RESULTS

### Dietary fibres have limited effects on gut microbial α-diversity

We sequenced 78 zebrafish intestinal samples (n = 10 per sex per diet) (refer to **Fig. S3**) after the three-week dietary intervention. Overall, Fusobacteriota and Pseudomonadota (Proteobacteria) were the top two phyla across our samples, with *Cetobacterium*, *Aeromonas*, and *Plesiomonas* being the core genera (**Fig. S4** and **Fig. S5**).

After running a single-shot rarefaction (depth = 52,938), we did not observe statistically significant changes in the α-diversity indices (Observed, FaithPD, Shannon, and Simpson) across the four diets (Control, 15AP, 15C, or 30APC) (Kruskal-Wallis for “Diet”, *p* > 0.05). We then reassessed the α-diversity indices by stratifying each diet by sex. Notably, we only detected a significant sex effect for the FaithPD index (*F*_1,55.017_ = 8.22, *p* = 0.006), but the diet effect for the FaithPD index was not statistically significant (*F*_3,12.352_ = 0.23, *p* = 0.874). We then compared α-diversity between females and males for the FaithPD index under each of the four diets. Under the combined 30APC diet, females showed lower FaithPD α-diversity than males (estimate = −12.32, *p* = 0.008) (**Fig. 1B**). In other words, under 30APC, male zebrafish had more diverse taxa than females at the phylogenetic level. We did not detect a statistically significant sex difference in the FaithPD α-diversity for the other three diets (Control, 15AP, 15C) (Fig. 1A, C, and **D**).

**Fig. 1:**
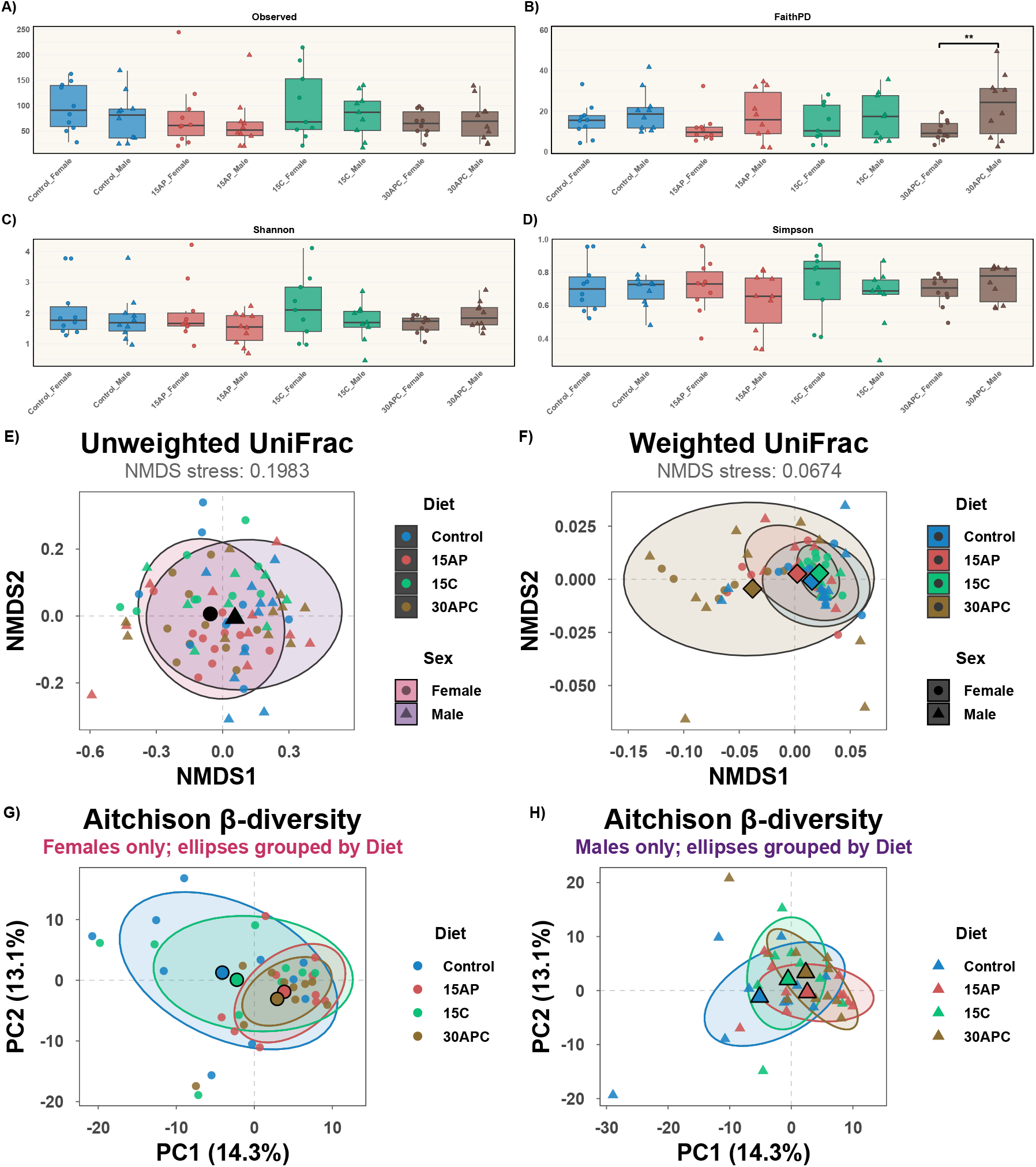
α- and β-diversity. (**A-D**) α-diversity indices box plots. (**E**) Unweighted phylogenetic-based β-diversity grouped by sex. (**F**) Weighted phylogenetic-based β-diversity grouped by diet. (**G**) Aitchison β-diversity based on CLR-transformed microbial composition, grouped by diet in females. (**H**) Aitchison β-diversity based on CLR-transformed microbial composition, grouped by diet in males. Each point represents an individual sample. Ordinations provide a reduced-dimensional representation of the underlying distance structure. Thus, interpret centroid proximity alongside the corresponding statistical analyses. ** denotes *p* < 0.01.

### Dietary fibres modulate the overall composition of the gut microbial β-diversity

#### UniFrac β-diversity

Using a phylogenetic approach, we then assessed the β-diversity among the samples from the four diets and both sexes. We found no significant “Diet:Sex” interaction for both the unweighted (*F* = 1.09, df = 3, R^2^ = 0.04, *p* = 0.226) and weighted UniFrac metrics (*F* = 0.55, df = 3, R^2^ = 0.02, *p* = 0.702). Notably, sex was the only significant factor associated with unweighted UniFrac β-diversity (*F* = 2.13, df = 1, R^2^ = 0.027, *p* = 0.001) (**Fig. 1E**). However, pairwise comparisons between females and males within each diet were not statistically significant (*p* > 0.05) (**Fig. S6**). Within each sex, microbial community composition differed between 15AP and 15C within females (*F* = 1.79, df = 1, R^2^ = 0.1, *p*_adj_ = 0.032). There was no dispersion by sex for unweighted UniFrac (*F* = 1.84, *p* = 0.172).

Next, we explored the abundance-weighted phylogenetic community structure using weighted UniFrac across the four diets. Notably, we found a significant “Diet” effect (*F* = 8.24, df = 3, R^2^ = 0.25, *p* < 0.001), but not “Sex” effect (*F* = 0.74, df = 1, R^2^ = 0.008, *p* = 0.417). As a post hoc analysis, we then conducted pairwise weighted UniFrac PERMANOVA comparing all six combinations. Interestingly, the microbial community composition in fish under the 30APC diet was significantly different compared to the Control (*F* = 10.515, R^2^ = 0.217, *p*_adj_ = 0.013) (**Fig. 1F**). Additionally, microbial community composition under 30APC differed significantly from both 15AP (*F* = 5.853, R^2^ = 0.133, *p*_adj_ = 0.04) and 15C (*F* = 14.185, R^2^ = 0.283, *p*_adj_ = 0.002). Thus, microbial diversity under 30APC differed from all three diets (**Fig. 1F**). In contrast, microbial communities did not differ between 15AP and the Control, or between 15C and the Control (both *p*_adj_ > 0.05). However, the weighted β-diversity differed significantly between 15AP and 15C (*F* = 5.952, R^2^ = 0.142, *p*_adj_ = 0.04) (**Fig. 1F**. We then performed sex-stratified follow-up analyses to assess these dietary contrasts separately in females and males. In females, the weighted β-diversity under 30APC was statistically different from that of Control (*F* = 12.03, R^2^ = 0.4, *p*_adj_ = 0.009) and 15C (*F* = 15.07, R^2^ = 0.47, *p*_adj_ = 0.005), but not 15AP (*F* = 3.84, R^2^ = 0.18, *p*_adj_ = 0.19). In males, none of the six diet pairwise comparisons reached statistical significance (all *p*_adj_ > 0.05 (**Fig. S7**).

After checking the homogeneity of dispersion for the weighted UniFrac metric, we found no dispersion by sex (*F* = 1.13, *p* = 0.3). However, microbial community dispersion differed significantly among the diets for weighted UniFrac (*F* = 11.48, *p* < 0.001), indicating that differences in within-group variability may have contributed to the PERMANOVA outcome. More specifically, microbial communities under 30APC were more dispersed than those under the Control, 15AP, and 15C diets (all *p*_adj_ < 0.001) (**Fig. 1F**). In follow-up analyses stratified by sex, the microbiome in male fish under 30APC was significantly more dispersed than that in male fish under the Control (*p*_adj_ = 0.02), 15AP (*p*_adj_ = 0.005), and 15C diets (*p*_adj_ < 0.001). In females, we detected significant dispersion only between 30APC and 15C (*p*_adj_ = 0.01). Overall, our data suggest that 30APC is associated with changes in both gut microbial community composition (weighted UniFrac) and inter-individual community variability (dispersion test).

#### Aitchison β-diversity

We then evaluated β-diversity using a compositional data analysis framework, applying Aitchison distance after a centred log-ratio (CLR) transformation to mitigate the confounding effects of heterogeneous dispersion observed with traditional distance metrics (UniFrac). Thus, we used the Aitchison distance as the primary measure of β-diversity in this study. As in the previous analysis, we did not detect a significant “Diet:Sex” interaction (*F* = 0.90, df = 3, R^2^ = 0.03, *p* = 0.694). Instead, we found a significant diet effect on microbiome composition (*F* = 2.566, df = 3, R^2^ = 0.094, *p* < 0.001), but no significant sex effect (*F* = 1.33, df = 1, R^2^ = 0.016, *p* = 0.124). Interestingly, unlike weighted β-diversity, Aitchison distance differed significantly in five of the six pairwise dietary comparisons (*p*_adj_ < 0.05), except between 15AP and 30APC, which was non-significant (*F* = 1.29, df = 1, R^2^ = 0.033, *p*_adj_ = 0.151). Also, after the follow-up analyses stratified by sex, the microbial community composition did not differ when contrasting 15AP vs 30APC in females (*F* = 0.89, df = 1, R^2^ = 0.047, *p*_adj_ = 0.611) (**Fig. 1G**), nor in males (*F* = 1.34, df = 1, R^2^ = 0.07, *p*_adj_ = 0.241) (**Fig. 1H**). Yet, in both sexes, the microbiome composition differed significantly between 15C and 30APC (females: *F* = 2.13, df = 1, R^2^ = 0.111, *p*_adj_ = 0.016; males: *F* = 2.19, df = 1, R^2^ = 0.114, *p*_adj_*=* 0.007) (**Fig. 1G-H**). No significant differences in dispersion were detected by “Diet” (*F* = 1.68, *p* = 0.179) or “Sex” (*F* = 0.12, *p* = 0.731) for the CLR/Aitchison distance β-diversity.

### Apple pectin and cellulose shape the microbial community through non-additive interactions

Having established that the diets influenced overall microbial community composition, we decomposed the diets into their underlying DF components to test whether the combined 30APC response could be explained by the additive effects of AP and C (AP + C), or whether their combination deviated from additivity (AP × C).

Using the previously calculated Aitchison distances, we conducted a sex-adjusted PERMANOVA with the apple pectin component and cellulose component as fixed effects, along with their interaction (AP component + C component + AP component:C component). We identified a significant main effect of the apple pectin component (*F* = 4.43, df = 1, R^2^ = 0.054, *p* < 0.001), but not the cellulose component (*F* = 1.32, df = 1, R^2^ = 0.016, *p* = 0.133). Interestingly, we detected a statistically significant interaction between the apple pectin and cellulose components (*F* = 1.95, df = 1, R^2^ = 0.024, *p* = 0.009), emphasising that the effect of each dietary component on microbial composition depends on the presence of the other. In other words, the significant interaction between apple pectin and cellulose highlights a non-additive effect of the two dietary components on the gut microbial composition (mathematically: effect (AP + C) ≠ effect (AP) + effect (C)). For statistical rigour, we also tested if this non-additive response differed by sex. Yet, we found no statistically significant effect for the “Sex:AP component:C Component” interaction (*F* = 0.69, df = 1, R^2^ = 0.009, *p* = 0.883) (**Fig. S8**).

Next, to further characterise the non-additive interaction and its direction, we compared the observed microbial composition of the combined 30APC diet with what we expected under an additive response. The observed 30APC community was closer to the Control centroid than predicted under an additive response (10.46 vs 17.52 Aitchison units). Also, the community composition under the combined diet (30APC) more closely resembled that induced by 15AP than 15C (Δ distance = −4.88, bootstrap 95% CI [-7.29 to −1.51]). The results indicate that the combined effect of apple pectin and cellulose components was less than additive (sub-additive) relative to the expected additive effect of apple pectin and cellulose individually (**Fig. 2**). The same directional pattern was observed when females and males were examined separately in the follow-up analyses.

**Fig. 2:**
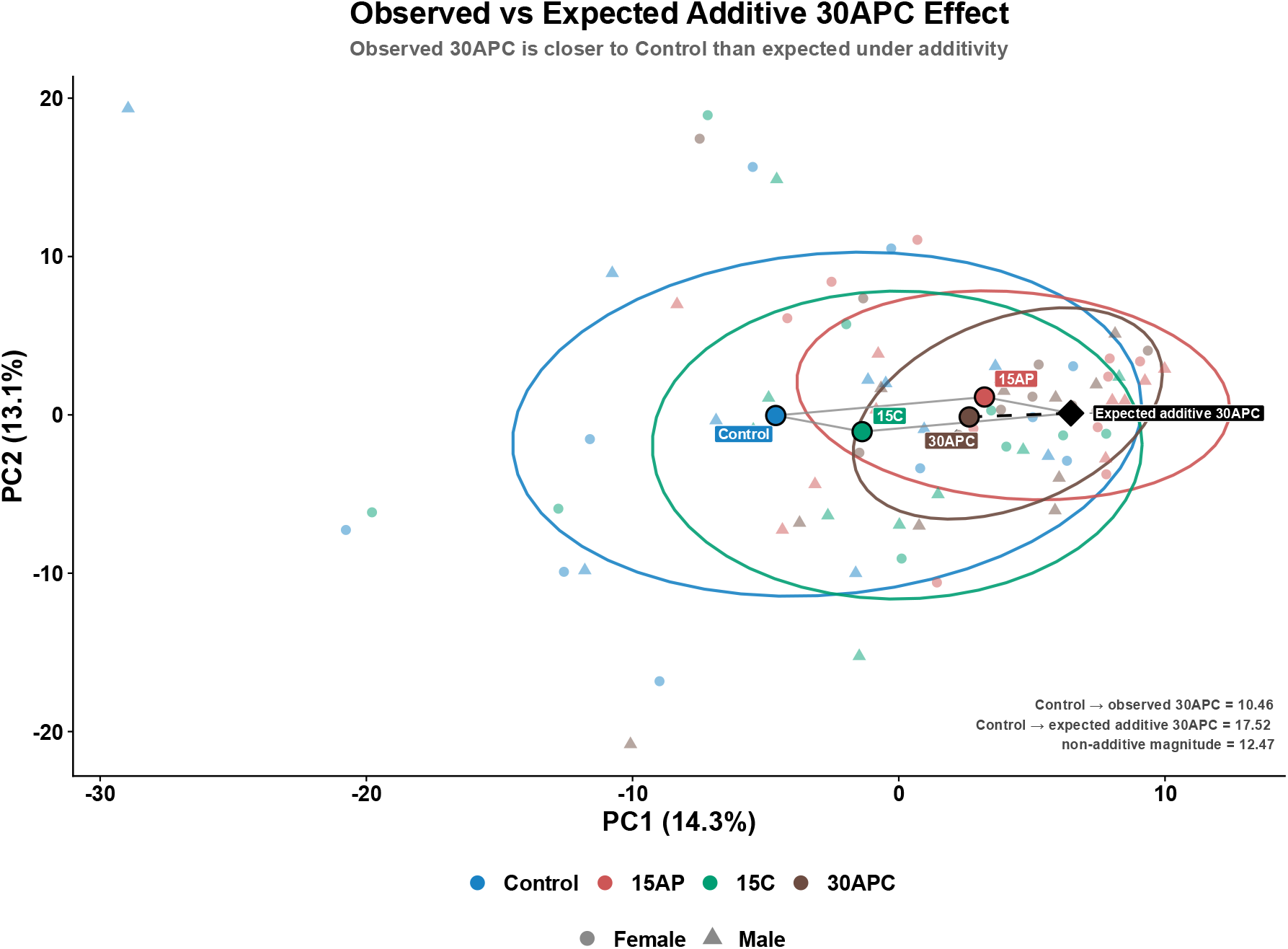
Non-additive effects on gut microbiome community composition. Centroid distances were calculated in the full Aitchison space. The observed 30APC centroid was closer to Control (10.46 Aitchison units) than the expected additive centroid (17.52 Aitchison units). The non-additive deviation (12.47 Aitchison units) represents the distance between the observed 30APC and the expected additive 30APC. The expected additive centroid was calculated as 15AP + 15C − Control.

### Dietary fibres shape the gut microbiome at the community level

We found that the gut microbiome responded differently across the diets. Next, we assessed whether these community-level differences extended to specific bacterial taxa, using targeted abundance comparisons followed by broader multigroup differential-abundance analysis with ANCOM-BC2.

At the genus level, across all four diets, we identified several differentially abundant taxa (primary analysis), which included *Plesiomonas* (*q* < 0.001), *Pseudomonas* (*q* = 0.001)*, Ensifer* (*q* = 0.005)*, Aquihabitans* (*q* = 0.03)*, unclassified_Hyphomicrobiaceae* (*q* = 0.03), and *Phreatobacter* (*q* = 0.04). However, only *Plesiomonas* was robust to the sensitivity analysis for pseudo-count handling. We then compared the top five genera in each experimental diet to the Control and found that, despite changes in abundance patterns across genera, statistical significance was not achieved (*q* > 0.05) (**Fig. S9**). Notably, we did not detect differentially abundant genera even after running pairwise contrasts of the experimental diets (**Fig. 3A**). We then repeated the analysis at the family level. Interestingly, although the experimental diets were not significantly different from the Control, Lachnospiraceae, a family notably associated with SCFA production^15^, was significantly more abundant in fish on the 15AP diet than in those on 15C (*q* = 0.003 and passed the sensitivity analysis) (**Fig. 3B**). Finally, the non-additive effect observed at the whole-community level was not attributable to significant AP × C interactions in individual bacterial genera or families after FDR correction (**Fig. S10** and **Fig. S11**). Overall, our results suggest diet-associated restructuring at the community level rather than selection for individual taxa.

**Fig. 3:**
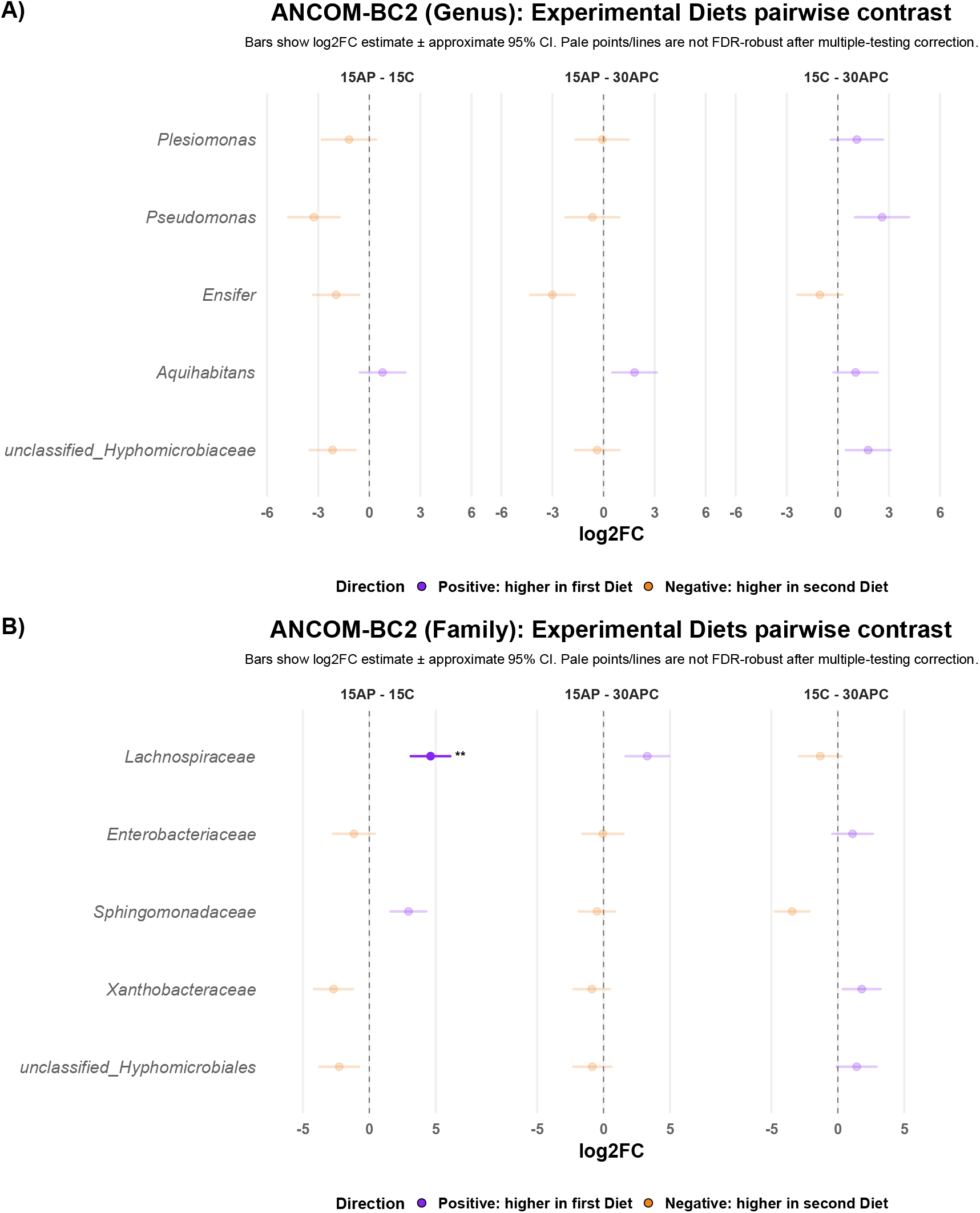
Differential abundance analysis comparing experimental diets. (A) ANCOM-BC2 at the genus level. (B) ANCOM-BC2 at the family level. Each point represents the model-estimated log2FC, and the horizontal error bar represents its 95% confidence interval. ** denotes *q* < 0.01

### Dietary fibres influence predicted microbial metabolic pathways

We used Phylogenetic Investigation of Communities by Reconstruction of Unobserved States (PICRUSt2) to investigate whether the taxonomic differences observed across diets were accompanied by differences in predicted microbial functional profile. We conducted the analysis using the Aitchison distances previously calculated for β-diversity. From our analysis, we only detected a statistically significant effect of “Diet” (*F* = 1.80, df = 3, R^2^ = 0.068, *p* = 0.033), but not of “Sex” (*F* = 0.74, df = 1, R^2^ = 0.009, *p* = 0.611) on the predicted functional composition. After establishing a diet-associated effect on the predicted functional composition, we explored individual metabolic pathways (https://biocyc.org/meta/) for their functional relevance to our study and the general effects of DFs on the gut microbiome. Notably, after BH (Benjamini-Hochberg) correction, we identified 16 diet-associated pathways related to cellular metabolism, fermentation, nucleotide biosynthesis, and cell wall synthesis (**Fig. 4A**). In our pairwise comparison between diets, we detected predicted functional differences in all five contrasts except the 15AP to 30APC contrast, suggesting that the microbial communities in those two diets may share similar predicted metabolic pathways (**Fig. 4B**). Finally, we tested whether the non-additive AP × C effect observed in Aitchison β-diversity was also reflected in the PICRUSt2-predicted functional profiles. In contrast to the gut microbial composition, the predicted functional profile under 30APC did not significantly deviate from additivity (*F* = 1.13, df = 1, R^2^ = 0.01, *p* = 0.287).

**Fig. 4:**
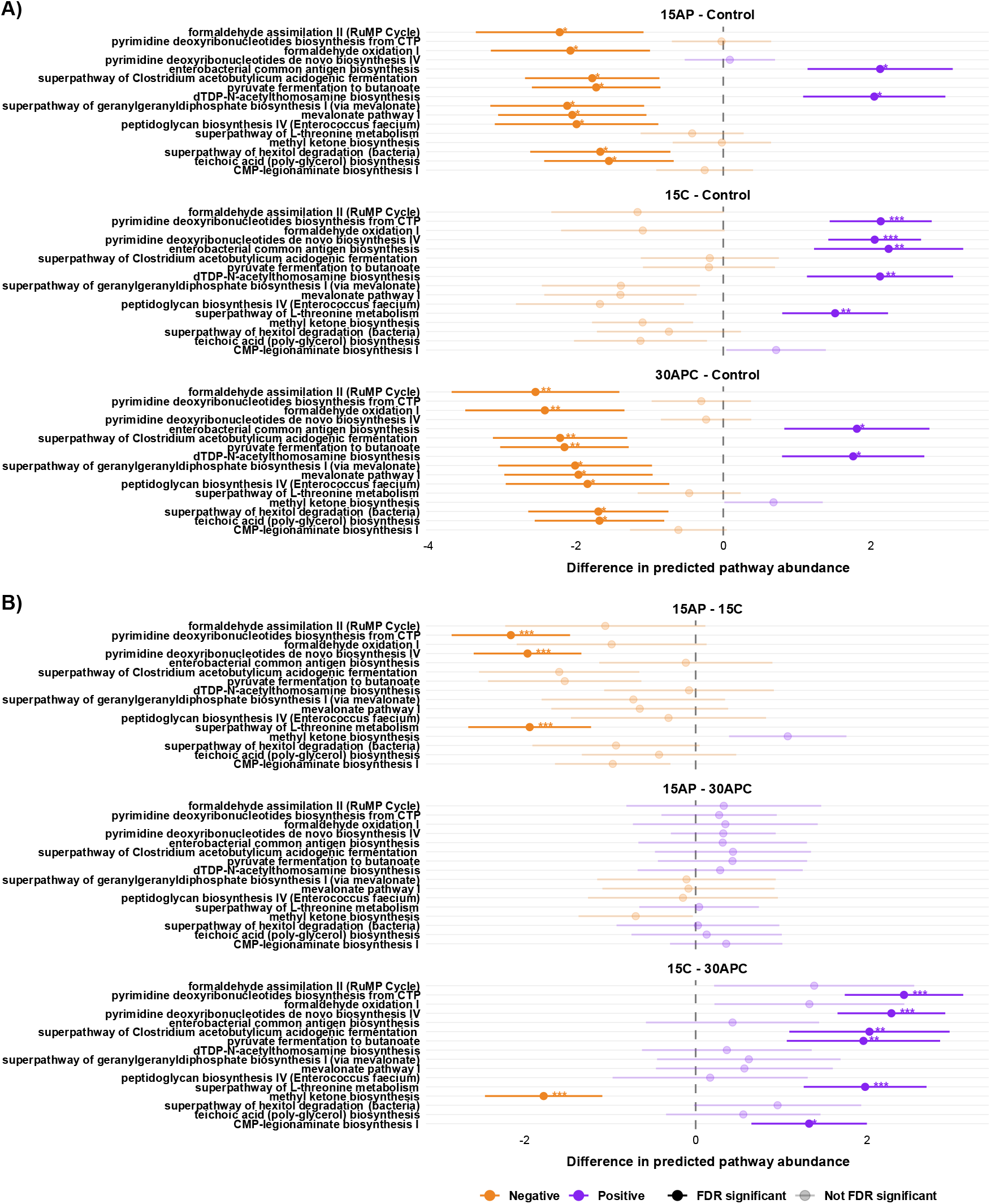
Predicted microbial metabolic pathways. (A) Experimental diet compared with Control. (B) Pairwise comparisons among experimental diets. Points represent model-estimated differences, with horizontal error bars indicating 95% confidence intervals. Positive values indicate greater predicted pathway abundance in the first diet of the contrast, whereas negative values indicate greater predicted abundance in the second diet. FDR adjusted * *p* < 0.05, ** *p* < 0.01, *** *p* < 0.001.

### Sex-specific tissue compartmentalisation of SCFAs

At the end of the three-week dietary intervention, we collected intestinal and brain samples to measure tissue SCFA levels across the gut-brain axis. Given that the different DFs altered gut microbial community composition and predicted microbial metabolic pathways, we next examined whether these changes were accompanied by corresponding differences in intestinal and brain SCFA availability.

In this study, we quantified six metabolites (acetate, propionate, isobutyrate, butyrate, valerate, and caproate) across both tissues. In the intestines of female and male fish, the three most abundant SCFAs were acetate, valerate, and propionate, but in different proportions. On average (n = 28), female intestines had 81% acetate, 14% valerate, and 4% propionate in the total SCFA pool. In male intestines, acetate accounted for 91% of the total SCFA pool, while propionate and valerate accounted for 5% and 2%, respectively. In the brain, acetate, propionate, and butyrate were the three most abundant SCFAs in both females and males, with acetate accounting for about 90% of the total brain SCFA profile.

The PCA (principal component analysis) showed clear separation of SCFAs by tissue type, indicating that the intestine and the brain function as distinct compartments in SCFA metabolism (**Fig. S12**). Additionally, we found that tissue SCFA availability differed between the sexes for all individual metabolites, with a significant “Sex:Tissue” interaction for each metabolite (lmer, *p* < 0.05). The full statistical model output for each metabolite is presented in the **Fig. S13**. Collectively, we found that SCFA pool availability across the gut-brain axis depends on the tissue and differs by sex. For acetate, the direction of sexual dimorphism reversed between brain and intestine. Specifically, females had lower intestinal acetate (estimate = −0.168, *p* < 0.001), but higher brain acetate (estimate = 0.078, *p* = 0.049) than males (**Fig. 5A**). More broadly, intestinal SCFA concentrations were significantly lower in females than in males for all measured metabolites, except valerate, which was significantly higher in the females than in the males (estimate = 0.418, *p* < 0.001) (**Fig. 5I**).

**Fig. 5:**
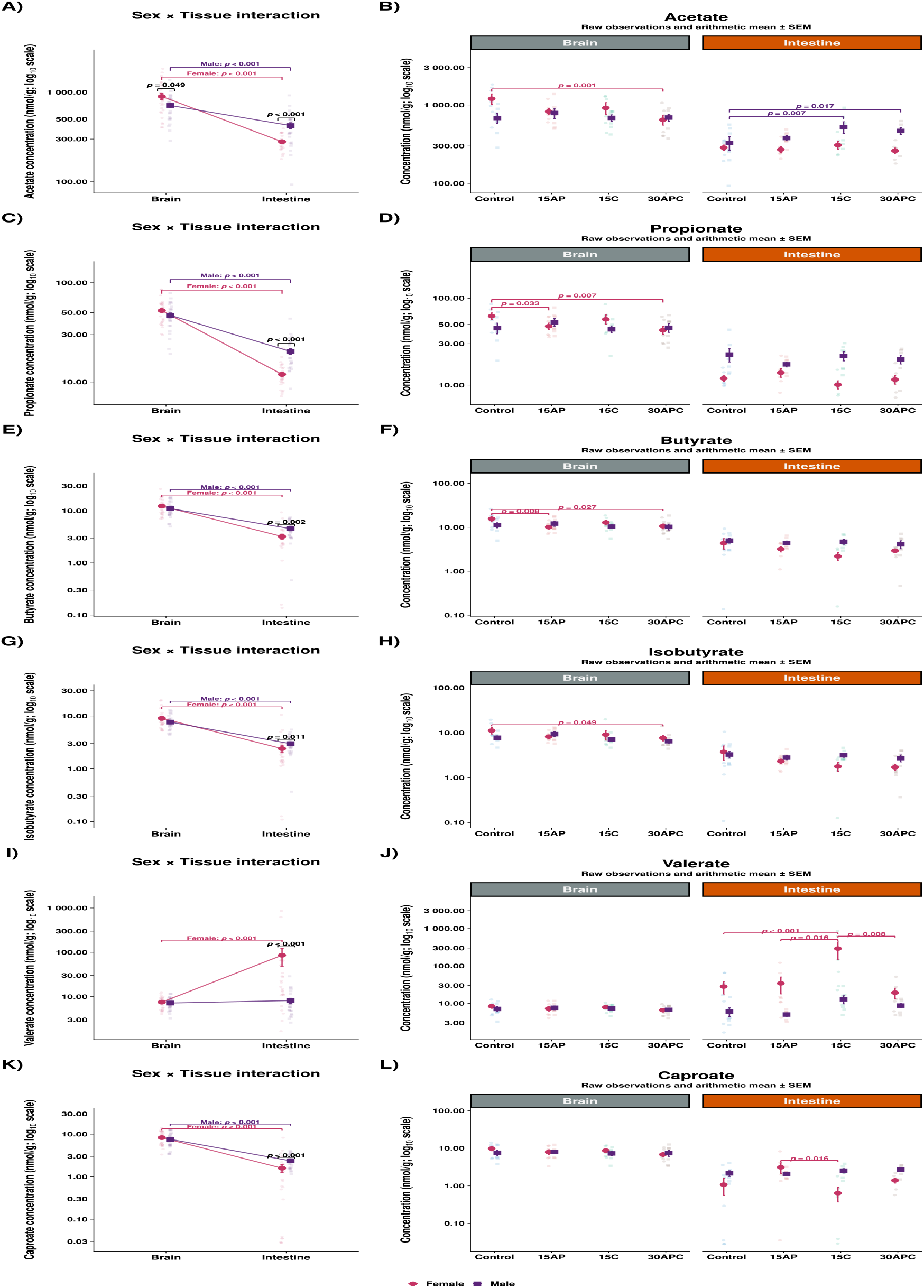
SCFAs concentrations across the gut-brain axis. (A-B) Acetate concentrations. (C-D) Propionate concentrations. (E-F) Butyrate concentrations. (G-H) Isobutyrate concentrations. (I-J) Valerate concentrations. (K-L) Caproate concentrations. SCFA concentrations are displayed on a log10-transformed y-axis for better visualisation. Dark points represent group means, overlaid on individual sample points, and error bars represent ± SEM.

### Dietary fibres effects on SCFA availability depend on sex and tissue

Having established sex- and tissue-dependent differences in SCFA availability, we explored the effects of the DFs. For all SCFAs concentrations, except butyrate and isobutyrate, dietary responses depended on sex and/or tissue (significant Diet:Sex, Diet:Tissue, or Diet:Sex:Tissue interaction) (refer to **Fig. S13** for full statistical models).

First, we examined the effect of “Diet” within the relevant “Sex:Tissue” combinations, comparing experimental diets with the Control. In male intestines, compared to Control, acetate was significantly higher under 15C (estimate = 0.257, *p*_adj_ = 0.007) and 30APC (estimate = 0.237, *p*_adj_ = 0.017) (**Fig. 5B**). On the other hand, in female intestines, we did not detect a significant change in acetate level across all pairwise diet comparisons (*p*_adj_ > 0.05) (**Fig. 5B**). Moreover, compared to Control, female intestinal valerate was significantly higher under 15C (estimate = 0.66, *p*_adj_ < 0.001) (**Fig. 5J**). In female brains, 15AP was associated with significantly lower propionate (estimate = −0.177, *p*_adj_ = 0.033) and butyrate (estimate = −1.796, *p*_adj_ = 0.008) than Control (Fig. 5D and **F**). Similarly, female 30APC fish had significantly lower brain concentrations of acetate, propionate, butyrate, and isobutyrate than female Control (all *p*_adj_ < 0.05) (**Fig. 5B, D, F** and **H**). No statistically significant differences were detected for any of the six SCFAs in male brains compared with the male Control (all *p*_adj_ > 0.05) (**Fig. 5B, D, F, H, J** and **L**).

For propionate and caproate, our results show that the diet effect was both tissue-dependent and differed between females and males (Diet:Sex:Tissue interaction: *F*_3,47.18_ = 4.3, *p* = 0.009 and *F*_3,88.4_ = 3.1, *p* = 0.031, respectively). For propionate, this was primarily reflected in significantly lower brain concentrations in females under 15AP (estimate = −0.177, *p*_adj_ = 0.033) and 30APC (estimate = −0.211, *p*_adj_ = 0.007) than in female Control. This pattern was not evident in the female intestine (**Fig. 5D**). For caproate, post-hoc contrasts did not identify a specific dietary comparison driving the significant three-way interaction (*p*_adj_ > 0.05) (**Fig. 5L**).

### Individual and combined dietary fibres induce distinct SCFA responses

Using the same fitted model for each SCFA, we next examined pairwise contrasts among the experimental diets. We first compared 15AP and 15C to characterise differences between the individual DFs, before comparing each with 30APC to assess whether the combined diet produced a distinct response.

In female intestines, valerate was significantly lower under 15AP than under 15C (estimate = −0.492, *p*_adj_ = 0.016), but not significantly different to 30APC (estimate = 0.04, *p*_adj_ = 0.993). Yet, intestinal valerate in female fish under 15C was significantly higher than 30APC (estimate = 0.53, *p*_adj_ = 0.008) (**Fig. 5J**). This suggests that the valerate intestinal combined phenotype (30APC) more closely resembled apple pectin (15AP) than cellulose (15C) in female fish. For caproate, intestinal levels were significantly higher in 15AP females than in 15C females (estimate = 1.5, *p*_adj_ = 0.016). However, female intestinal caproate under 15AP or 15C did not differ statistically compared to 30APC (estimate = 0.66, *p*_adj_ = 0.503 and estimate = −0.84, *p*_adj_ = 0.324, respectively) (**Fig. 5L**). For propionate, the 15AP vs 15C comparison differed significantly across sex and tissue (*p*_adj_ = 0.022), consistent with a crossover pattern (opposite direction depending on the tissue). However, the corresponding within-tissue comparisons were not statistically significant (*p*_adj_ > 0.05) (**Fig. 5D**). Finally, we found no evidence that the SCFA response deviated from additivity, as indicated by the lack of a significant AP × C interaction for any of the six measured SCFAs (**Fig. S14**).

### Sex significantly determines liver metabolic variation

In this study, we broadened our metabolomics analysis to explore a wide range of liver metabolites, thereby gaining a more comprehensive understanding of diet-host interactions. We quantified 61 metabolites using both HILIC (Hydrophilic Interaction Liquid Chromatography) and reverse-phase chromatography (**Fig. 6**). The liver metabolomic pool included amino acids, amines, purines, acylcarnitines, and tryptophan. Collectively, these metabolite classes provide broad coverage of central energy, amino acid, nucleotide, and host-microbial metabolic pathways.

**Fig. 6:**
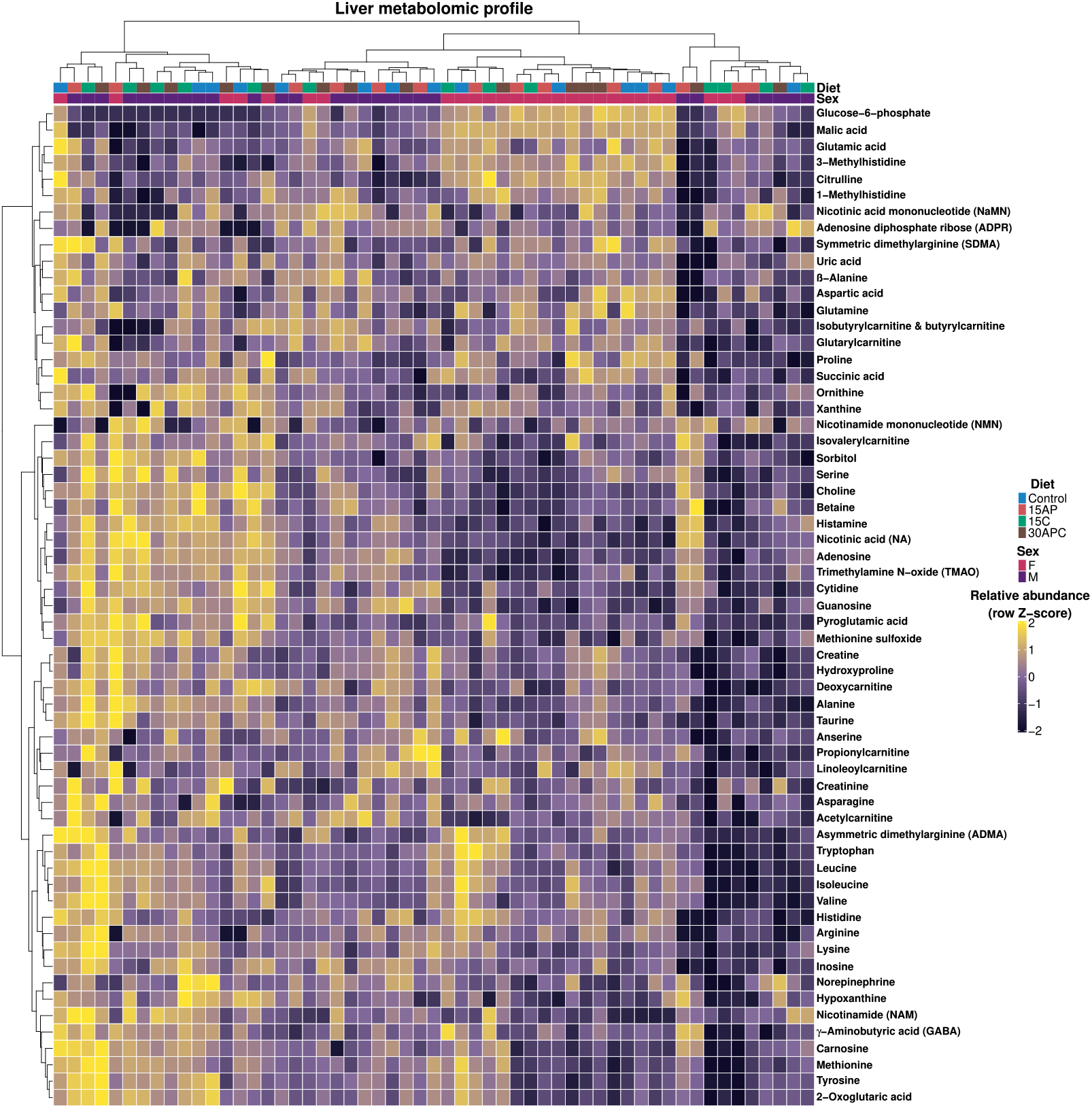
Liver metabolomic profile. Unsupervised hierarchical clustering heatmap showing metabolite abundance patterns across samples.

The liver metabolomic profiling heatmap suggests pronounced clustering by sex rather than diet (**Fig. 6**). Next, we statistically tested whether overall liver metabolite differed by “Diet”, “Sex”, and their interaction. Notably, only “Sex” significantly explained overall liver metabolic variation (*F* = 6.31, df = 1, R^2^ = 0.106, *p* < 0.001), with no “Diet” (*F* = 1.04, df = 3, R^2^ = 0.052, *p* = 0.396) or “Diet:Sex” effect (*F* = 1.01, df = 3, R^2^ = 0.051, *p* = 0.433). Moreover, we found no difference in dispersion between the sexes (*F* = 0.38, *p* = 0.552), which supports the multivariate metabolic analysis.

Next, we identified which metabolites, if any, differed significantly between females and males in the overall pool (averaged across all diets). After BH (Benjamini-Hochberg) false discovery rate correction, 20 of the 61 liver metabolites remain significantly associated with “Sex” (*p* < 0.05). **Table 1** lists 25 of the 61 liver metabolites that showed an unadjusted *p* < 0.05, of which 20 remained significant after FDR correction. It also indicates the effect and direction (F:M ratio > 1 = higher in females and vice versa). Overall, we identified sex as the primary driver of liver metabolite availability.

**Table 1:**
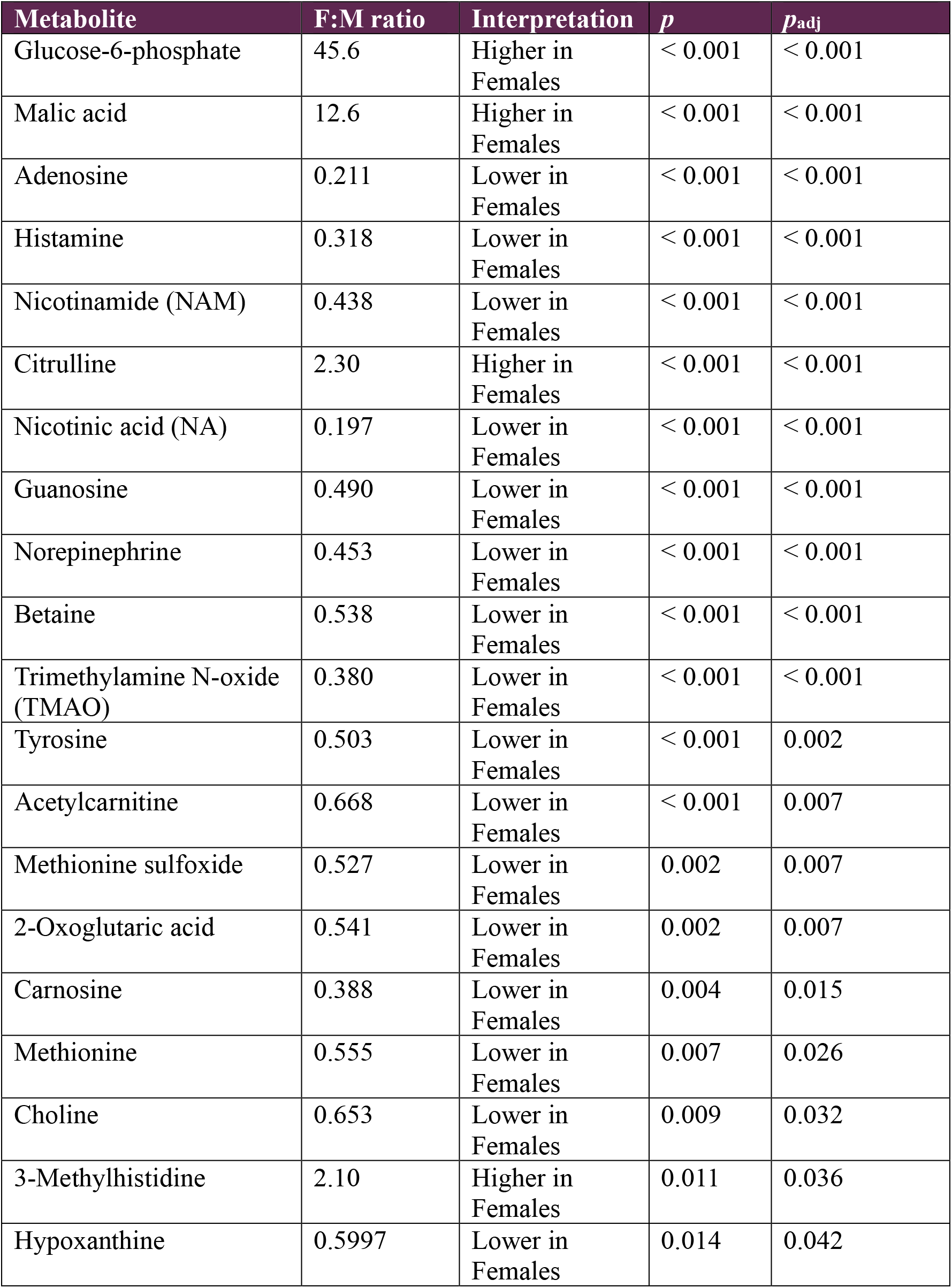
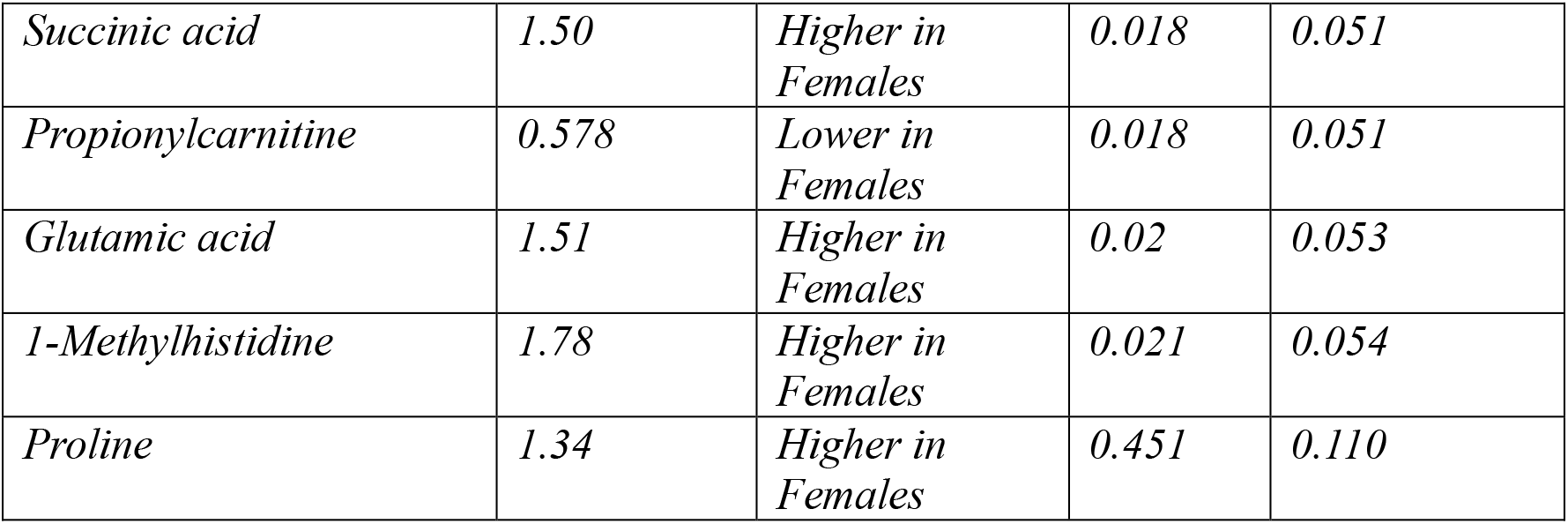
Liver metabolites differing significantly between sexes.

### Associations between predicted microbial functions and quantified tissue SCFAs

#### Predicted microbial metabolic pathways correspond with intestinal SCFAs in a metabolite-and pathway-dependent manner

We investigated whether the predicted microbial capacities inferred from our microbiome data (PICRUSt2) correlated with the quantified LC-MS/MS metabolites. We focused on the three major SCFAs (acetate, propionate, and butyrate), for which PICRUSt2 represents well-characterised microbial fermentation pathways. We excluded isobutyrate, valerate, and caproate because their production can involve more complex metabolic routes, including amino-acid fermentation and microbial chain elongation^16^, making the dataset’s predicted metabolic pathway proxies less reliable. Importantly, we conducted correlation analyses at the sex-stratified group level rather than the individual-fish level, as the microbiome and metabolomics datasets came from different, unmatched fish.

We first ran exploratory Spearman correlations (n = 8 group level) between intestinal concentrations of acetate, propionate, and butyrate and the selected predicted pathways, but found weak effect sizes (ρ) that did not reach statistical significance (*p* > 0.05) (**Fig. S15**). This indicates limited correlation between predicted microbial functions and quantified tissue SCFAs at the group level. Next, we assessed whether the dietary response in predicted microbial function was directionally concordant with that of the corresponding intestinal SCFA. For each sex, responses to 15AP, 15C, and 30APC were expressed relative to the Control (n = 6 diet-response observations). We summarised diet-response concordance using a four-quadrant plot (**Fig. S16**), where the upper-right and lower-left quadrants indicate directionally concordant increases and decreases, respectively. This analysis reflects only the direction of the SCFA response. Statistical significance should be determined from the adjusted contrasts in the main models.

For acetate, directional concordance was low for both pathway modules A and B (33% concordance; 2 of 6 samples). Module A consistently decreased across all diet responses, yet four samples had higher intestinal acetate than the Control (**Fig. 7A** and **Fig. 8**). Module B also showed mixed pathway responses that did not match the changes in intestinal acetate (**Fig. 7A** and **Fig. 8**). For propionate, we observed a 50:50 split between concordance and discordance in both the primary propionate pathway and the three-pathway composite (**Fig. 7B** and **Fig. 8**). Notably, for 15AP, intestinal propionate increased relative to the Control in females but decreased in males, even though both sexes showed the same lower predicted microbial function pattern. This finding further shows limited correspondence between the predicted microbial response and the quantified propionate intestinal response. For butyrate, directional concordance was low for both P163-PWY and PWY-5676 (33% concordance; 2 of 6 samples) (**Fig. 7C** and **Fig. 8**). Interestingly, for the butyrate CENTFERM-PWY pathway, we observed complete concordance (100%; 6 of 6 samples). Specifically, the dietary effects on the predicted pyruvate-to-butanoate fermentation pathway (CENTFERM-PWY) matched the lower quantified intestinal butyrate levels (**Fig. 7C** and **Fig. 8**). Notably, this observation aligns with our previous PICRUSt2 analysis, in which 15AP and 30APC were associated with significantly lower predicted representation of the pyruvate-to-butanoate fermentation pathway than Control (*p*_adj_ < 0.05) (**Fig. 4B** and **Fig. 8**). Ultimately, we observed a metabolite- and pathway-dependent pattern between the predicted microbial function and intestinal SCFAs. For completeness, we also present the same direction-concordance analysis comparing the experimental diets (**Fig. S17**).

**Fig. 7:**
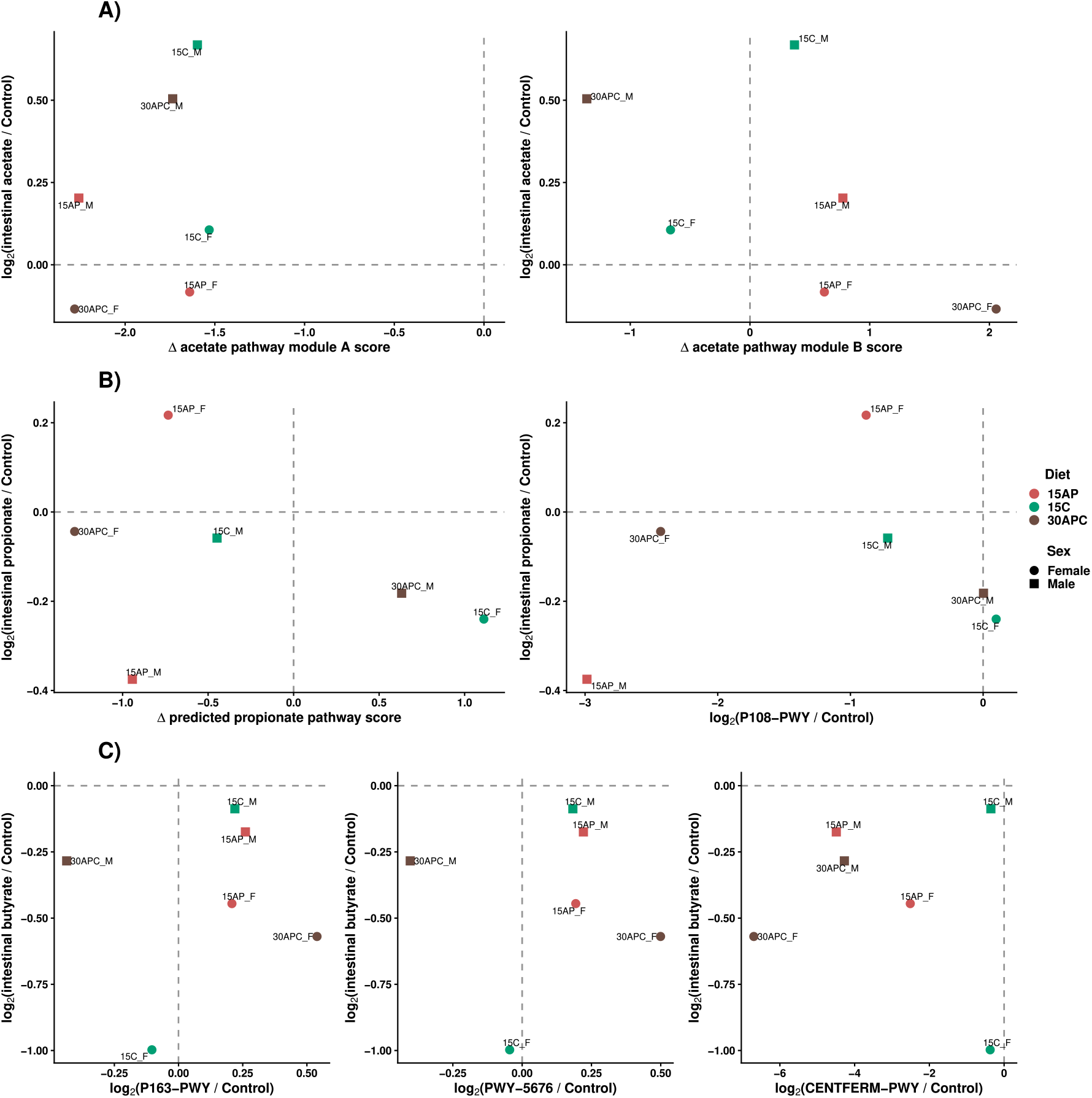
Direction-concordance analysis comparing experimental diets with the sex-matched Control in the intestine. The analysis compares the direction of change in quantified SCFA concentration with the direction of change in predicted microbial pathways. Concordant responses indicate that the SCFA and pathway changed in the same direction relative to Control, whereas discordant responses indicate changes in opposite directions. (**A**) Intestinal acetate. (**B**) Intestinal propionate. (**C**) Intestinal butyrate. For each axis, values are expressed as log2(experimental diet/Control); therefore, positive values indicate an increase relative to Control and negative values indicate a decrease. The analysis describes directional agreement only and not statistical inference.

**Fig. 8:**
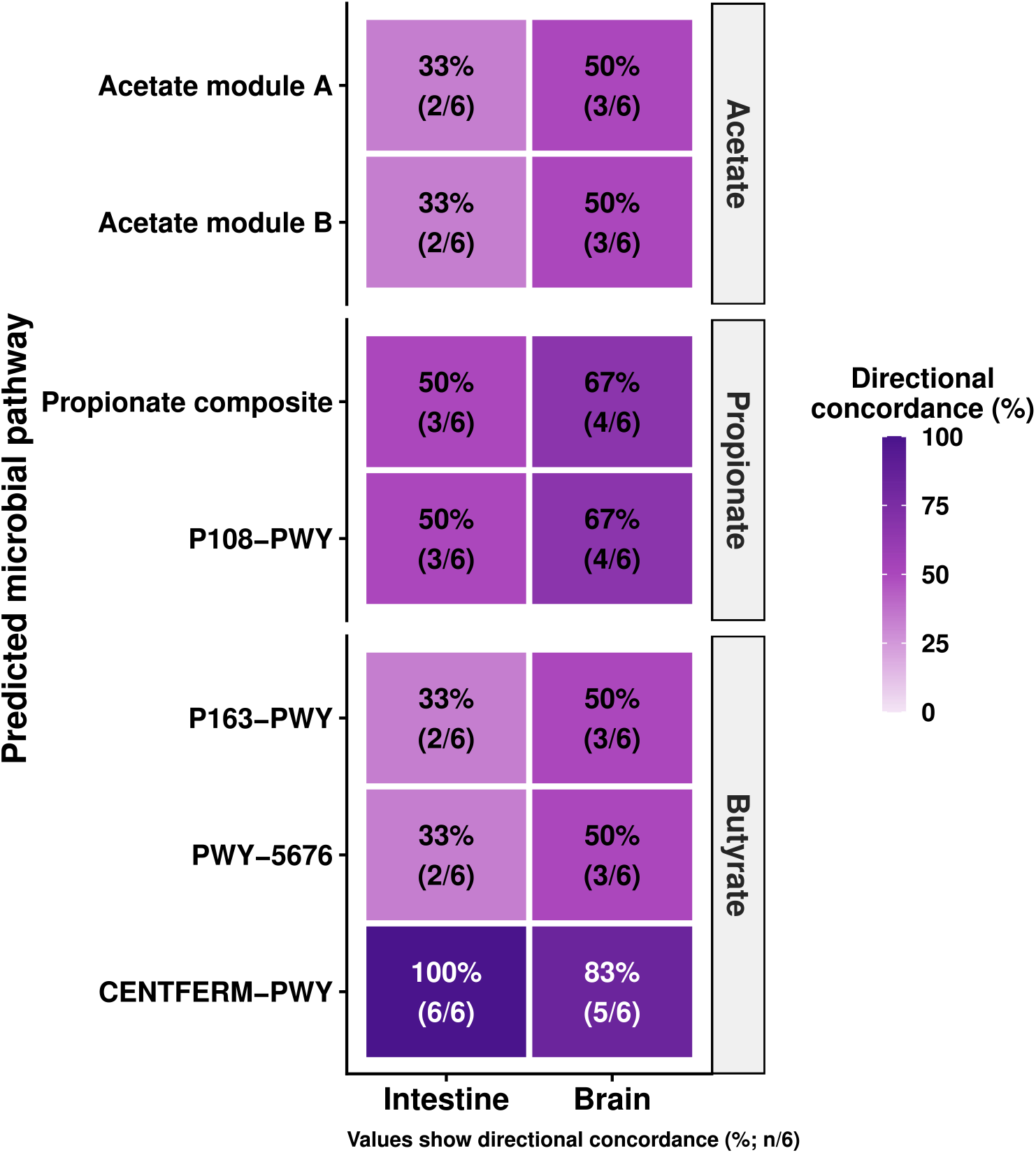
Summary of directional concordance between predicted microbial pathways and tissue SCFA concentrations. The percentage indicates the proportion of six sex-stratified comparisons between the experimental diet and control that show directional concordance in the intestine and the brain. (n = 6; three diet comparisons × two sexes).

#### The relationship between predicted microbial functions and SCFA availability differs along the gut-brain axis

We extended our analysis to test whether predicted gut microbial metabolic potential corresponded with SCFA availability beyond the intestine, specifically in the brain. Using the same pathways as in the previous analysis, we examined whether the direction of diet-associated changes in brain SCFAs relative to Control aligned with predicted gut microbial metabolic responses.

Similar to the intestinal exploratory Spearman correlations (n = 8 group level), we did not detect statistically significant correlations (*p* > 0.05) between brain acetate, propionate, and butyrate and the predicted microbial pathways. However, some brain comparisons showed larger Spearman correlation coefficients (ρ) than those observed in the intestine (**Fig. S15** and **Fig. S18**). We then compared the brain patterns with those observed in the intestine. The directional correspondence between predicted microbial functional responses and SCFA availability differed between the two tissues. While some pathway-metabolite relationships were preserved in the brain, others diverged from the intestinal response, suggesting that the correspondence between predicted microbial functional potential and tissue SCFA availability is not consistently aligned along the gut-brain axis (**Fig. 8** and **Fig. S19**).

### Post-intervention intestinal barrier gene expression

#### Sex-dependent intestinal gene expression varies in magnitude and direction

For the second phase of this study, we tested the expression of eight genes related to different aspects of the intestinal immune barrier response (**Fig. S20** and **Table S1**). Across all treatments, female intestinal tissues showed lower gene expression than male tissues (*p* < 0.05), except for *il1b* and *il6*, which did not differ significantly between the sexes. Interestingly, we detected a significant “Treatment:Sex” interaction on intestinal *hmgcra* expression (*F*_4,86_ = 3.92, *p* = 0.006). Also, intestinal *hmgcra* expression was significantly higher in females than males (log2FC = 1.84, *p* < 0.0001) (**Fig. 9**).

**Fig. 9:**
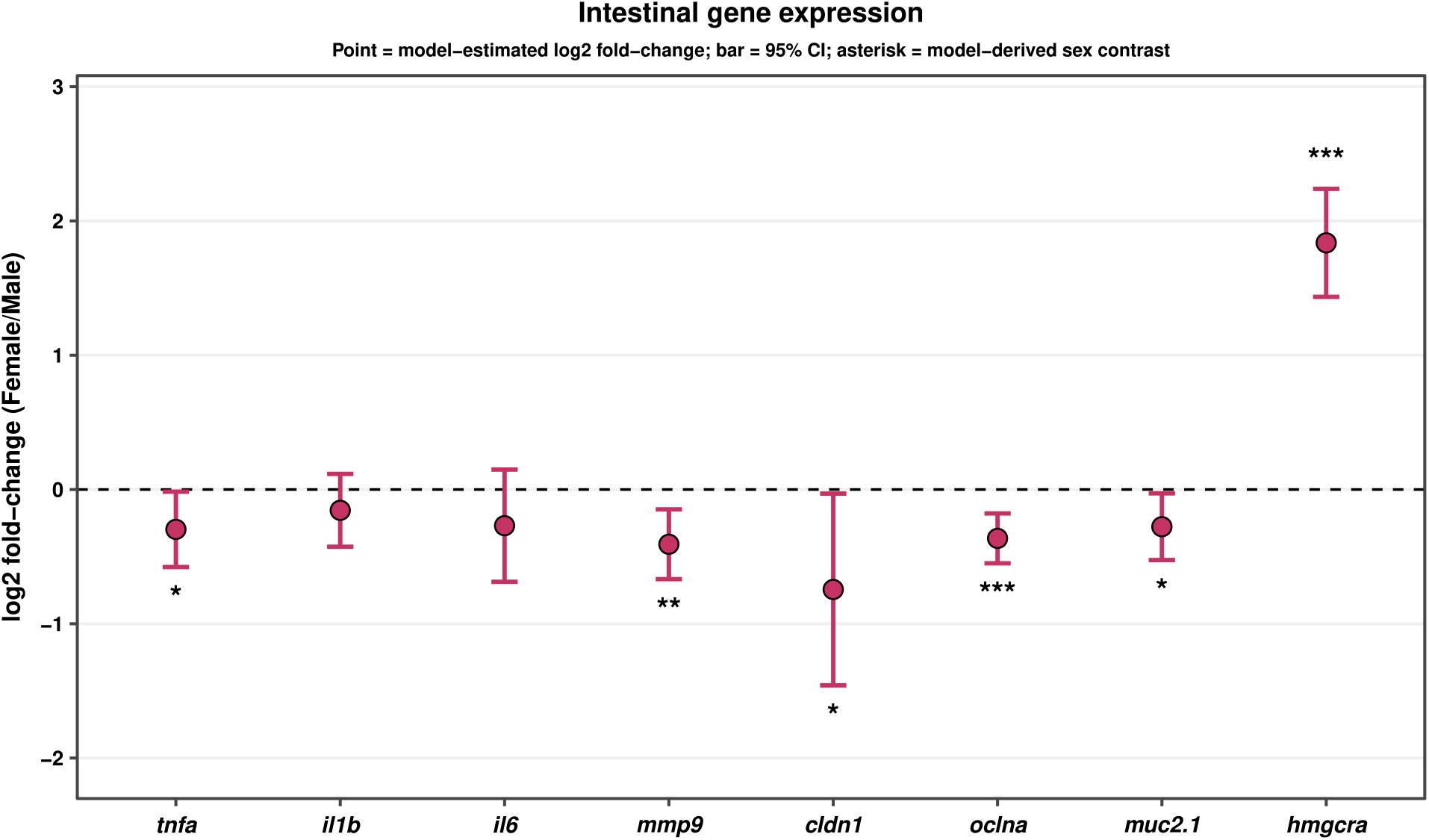
Intestinal gene expression in females relative to males. Each point is the model-estimated log2FC, and the vertical error bar is its 95% confidence interval. * *p* < 0.05, ** *p* < 0.01, and *** *p* < 0.001.

#### Prior dietary fibre intake modulates saponin-induced intestinal transcriptional responses, with significant effects detected in males

A key objective of this study was to investigate whether prior DF intake could attenuate the intestinal response to saponin-induced stress. Thus, we compared gene expression in Control fish exposed to saponin with that in the unchallenged Control group in both sexes. Notably, immune/inflammatory gene expression (*tnfa*, *il1b*, and *il6*) did not differ significantly between the two treatments (**Fig. S21A-C**). Moreover, *mmp9* expression, commonly upregulated during inflammatory tissue injury and pathological tissue remodelling^17^, also did not change after saponin challenge (**Fig. S21D**). We then examined tight junction genes (*cldn1* and *oclna*). There was no change in gene expression for *cldn1* in either sex. However, in male fish only, saponin upregulated *oclna* expression relative to the unchallenged Control group. Yet, this pattern was marginally significant (log2FC = 0.65, *p*_adj_ = 0.051) (**Fig. 10A** and **Fig. S21E**). In contrast, saponin exposure significantly upregulated both intestinal *muc2.1* (log2FC = 0.91, *p*_adj_ = 0.009) and *hmgcrca* (log2FC = 2.60, *p*_adj_ < 0.0001) compared to the unchallenged Control. Notably, this trend occurred only in male fish, not in females (Fig. 10B and **C**). Collectively, the results indicate that saponin administration induced detectable changes in genes related to mucus secretion^18^, cholesterol synthesis^19^, and possibly tight junction function, but only in males.

**Fig. 10:**
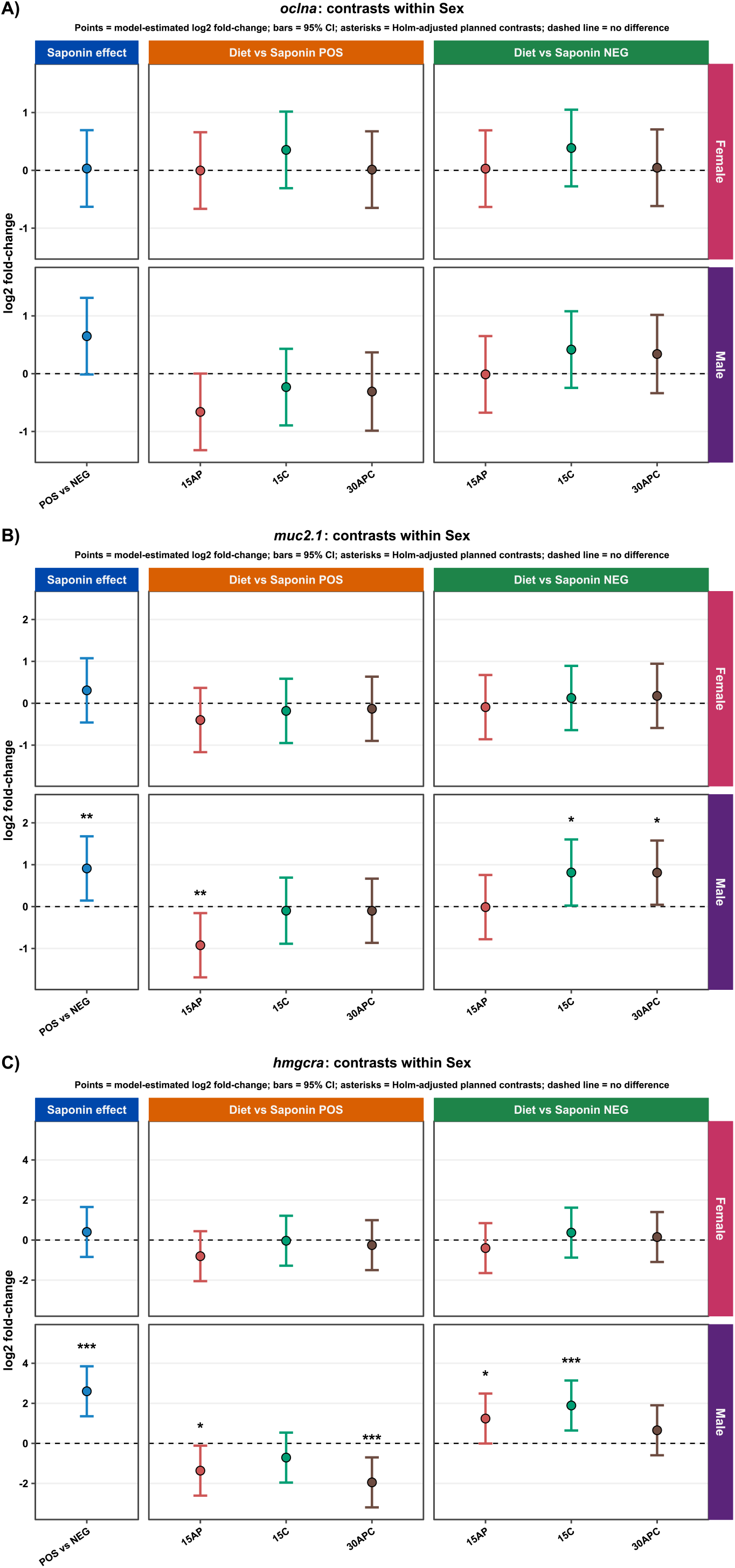
Intestinal *oclna*, *muc2.1*, and *hmgcra* gene expression across treatment comparisons. (**A**) Intestinal *oclna* gene expression. (**B**) Intestinal *muc2.1* gene expression. (**C**) Intestinal *hmgcra* gene expression. Each point is the model-estimated log2FC, and the vertical error bar is its 95% confidence interval. * *p* < 0.05, ** *p* < 0.01, and *** *p* < 0.001. Saponin POS refers to saponin-challenged Control, and Saponin NEG refers to unchallenged Control.

After establishing saponin’s effect in this experiment, we explored whether prior DF intake could alter post-saponin gene expression, focusing on *oclna*, *muc2.1*, and *hmgcra*. For this analysis, we compared gene expression in saponin-challenged fish previously fed 15AP, 15C, or 30APC with the saponin-challenged Control group. Interestingly, we found that male fish previously fed 15AP had significantly lower intestinal expression of *muc2.1* (log2FC = −0.92, *p*_adj_ = 0.009) and *hmgcra* (log2FC = −1.36, *p*_adj_ = 0.014) (Fig. 10B and **C**). Male *oclna* expression was also lower but marginally significant (log2FC = −0.66, *p*_adj_ = 0.051). Notably, fish previously on the 15C diet showed no difference in the expression of these three genes in either sex. However, male fish previously on the 30APC diet also showed significantly lower intestinal *hmgcra* expression (log2FC = −1.95, *p*_adj_ = 0.0003) than saponin-challenged Control fish (**Fig. 10C**).

We then compared the experimental dietary groups with saponin-negative Control fish (Control fish not exposed to saponin) to determine how gene expression following the saponin challenge compared with the unchallenged physiological baseline. Male fish from the 15AP diet showed *muc2.1* levels similar to the unchallenged Control group (log2FC = −0.013, *p*_adj_ = 1), consistent with expression levels being near the unchallenged baseline. Yet, male intestinal *muc2.1* expression was higher than the unchallenged baseline in fish previously fed either the 15C diet (log2FC = 0.81, *p*_adj_ = 0.02) or the 30APC diet (log2FC = 0.81, *p*_adj_ = 0.02), with comparable effects. In contrast, intestinal *hmgcra* gene expression was significantly higher in males from the 15AP (log2FC = 1.24, *p*_adj_ = 0.02) and 15C (log2FC = 1.89, *p*_adj_ < 0.001) compared to unchallenged Control (Fig. 10B and **C**). The results suggest that prior consumption of apple pectin-containing diets (15AP or 30APC) can attenuate saponin-induced intestinal transcriptional changes in male fish, with some gene expression remaining comparable to the unchallenged baseline. Consistent with this, post-saponin *muc2.1* expression differed between the experimental diets, with 15AP males differing significantly lower from both 15C (log2FC = −0.83, *p*_adj_ = 0.039) and 30APC (log2FC = −0.82, *p*_adj_ = 0.032). In contrast, 15C and 30APC fish showed comparable intestinal *muc2.1* responses (log2FC = 0.002, *p*_adj_ = 1.00) (**Fig. S22**). This pattern was not apparent for male *hmgcra* expression (**Fig. S23**).

## DISCUSSION

The health impact of DFs cannot be understood and predicted by considering their heterogeneous physicochemical properties in isolation^6,20,21^. Instead, their effects emerge from interactions with other dietary components, the microbiome, the host, and ultimately the crosstalk between all these elements. As shown in our previous work^3,22^, the host and its associated gut microbiome composition are key prerequisites shaping the biological context of DF responses. Here, we extend our investigation to show that DFs can interact within the same diet to produce a non-additive shift in the microbial community, which was not apparent at the predicted functional capacity level. We also explored the host microbiome-metabolism crosstalk by contrasting microbial responses with host tissue metabolite availability. Notably, our results indicate a lack of consistent concordance between the two. Despite that, the effects of prior DF exposure on the intestinal barrier seem to persist beyond active exposure.

Apple pectin (AP) and cellulose (C) are structurally and functionally distinct DFs that differ in solubility, structural complexity, and fermentability, all of which affect their accessibility to members of the gut microbiome^23,24^. In this study, microbial communities differed significantly between fish fed 15AP and 15C under both weighted UniFrac and Aitchison analyses. However, gut microbial community differences between 15AP and 30APC depended on the β-diversity metric. Specifically, these diets differed significantly under weighted UniFrac (phylogenetic structure), but not under Aitchison distance (**Fig. 1**). This suggests that AP in isolation can induce a phylogenetically distant community compared to when in combination with C. Notably, we also found that the microbiome in the 30APC fish was significantly more dispersed and phylogenetically hypervariable compared to the microbial communities in the other three diets: Control, 15AP, and 15C. This suggests that the microbial community under 30APC had reduced stability or greater heterogeneity in response to the more complex dietary environment. This greater dispersion might indicate the emergence of distinct responder vs non-responder phenotypes, whereby baseline differences in microbial community composition can contribute to greater inter-individual variability in response to increased substrate heterogeneity^5,25,26^. However, substrate complexity does not always result in a highly variable response. For example, increasing carbohydrate structural complexity, rather than variations in dietary substrates, could lead to a more consistent microbial community assembly^27^. The increased heterogeneity observed under 30APC suggested a possible interaction between AP and C within the microbial community. Consistent with this, the combined diet produced a less-than-additive shift in community composition, deviating from the response expected from the individual effects of AP and C. Mechanistically, each dietary component in the 30APC diet may exert distinct selective pressure on the microbial community while also modifying the community context in which the other component is utilised. We propose several mechanisms, which are not mutually exclusive, that could help explain the microbial response. AP is a relatively fermentable DF that can influence digesta viscosity, whereas C is less accessible to microbial fermentation. Their contrasting physicochemical properties may therefore jointly alter the gut’s metabolic and structural environment, with downstream effects on the microbial community^28^. Moreover, as the microbiome shifts in response to diet, different members of the microbial community may compete for available substrates, meaning that the selective pressures induced by the combined DFs may differ from those generated by either DF in isolation^5,29^. On the other hand, cross-feeding can occur when primary degraders break down a specific DF into products accessible to secondary degraders, while also generating metabolites that can serve as substrates for other community members^30^. Thus, adding an additional substrate may not drastically alter microbial composition if it supports overlapping microbial functions or is used by taxa already present in the community. Typically, a greater-than-additive response indicates synergy^5^. Although we strategically combined two DFs with distinct structural and physicochemical properties, their combination did not produce a synergistic response. We hypothesise that this less-than-additive response may partly reflect the specific components selected for the combined diet. For example, combining AP with a DF with different physicochemical and fermentative properties, such as the oat β-glucan used in our previous experiment^3^, could have produced a different or potentially stronger shift in the gut microbial community. However, this remains speculative, as combining two or more fermentable DFs does not necessarily produce a synergistic response. Studies suggest that non-additivity, and specifically true synergistic effects, may be more likely to arise when each dietary component does not favour direct microbial competition but instead caters more strongly to distinct functional and metabolic niches, potentially reducing functional redundancy and encouraging cross-feeding^5,31,32^. Importantly, not all DF combinations show such more-than-additive/synergism, with other mixed-substrate interventions instead producing additive responses^33^. Thus, the overall response depends on the existing microbial community, the degree of ecological overlap, and the capacity of the DFs to exert distinct selective pressures^29^.

We also explored the microbiome-metabolism crosstalk and found that predicted microbial pathways do not necessarily translate into tissue SCFA availability across the gut-brain axis (**Fig. 5** and **Fig. 8**). Based on our previous metabolomics data^3^ and further supported by this study, we infer that targeted metabolite quantification measures the concentration remaining in host tissue at the time of sampling, rather than the total amount produced or the extent of *in vivo* exposure. In other words, intestinal SCFAs produced by the gut microbiome through DF metabolism may be further utilised by members of the microbial community^30^, taken up and metabolised by intestinal epithelial cells in a metabolite-specific manner, or enter the portal circulation, where they undergo further hepatic metabolism before a proportion reaches the peripheral circulation and potentially the brain^34^. This long cascade of events, from SCFA synthesis in the gut to their eventual availability in distal tissues such as the brain, means that microbial production or predicted microbial capacity does not necessarily correspond quantitatively to the concentration measured in a particular tissue. Thus, to establish metabolic kinetics and flux along the gut-brain axis, we suggest sampling at several time points or using isotope tracing^35^. Despite this, we still detected changes in intestinal and brain SCFAs in response to the various DFs, influenced by interacting host factors, namely sex and tissue type (**Fig. 5**). Our results show that SCFA concentrations across tissues do not necessarily change in parallel. Evidence across species indicates that females and males differ in the regulation of energy metabolism and substrate utilisation across multiple tissues^36–38^. SCFAs produced by the gut microbiome are not metabolically inert^34^. For example, intestinal epithelial cells largely utilise butyrate, whereas acetate and propionate more readily enter the portal circulation. The liver then extensively metabolises propionate, while a greater proportion of acetate reaches the systemic circulation. In our study, lower brain SCFA availability cannot necessarily be interpreted as reduced brain exposure, as it could also reflect differences in uptake or utilisation within the brain, which may have led to the observed lower tissue concentration at the sampling time point. As zebrafish metabolomics advances, confirming these possibilities will require direct measurement of metabolic flux, which will most likely also differ by sex. We also confirm that sex is a major source of metabolic variation across multiple tissues, such as the intestine, brain, and even liver.

We have previously shown^3^ that the feeding pattern of DF consumption (regular vs intermittent) can also shape the host’s response. Nutritional recommendations support regular consumption of DFs from a diverse range of whole-food sources to promote health and reduce the risk of chronic disease^14^. However, it remains unclear whether the putative benefits of regular DF intake persist after cessation, whether the host reverts to baseline, or whether it leaves a physiological memory^39,40^. In this study, we tested this hypothesis by challenging the fish previously fed DFs with saponin. We aimed to induce intestinal inflammation, which could lead to intestinal barrier damage^41^. From our qPCR data, we did not detect signs of acute intestinal inflammation at the genetic level (*tnfa, il1b, il6,* and *mmp9*) or significant changes in core tight junction transmembrane components (*cldn1* and *oclna*). We conducted the saponin challenge daily for two weeks (14 days), a duration at which previous adult zebrafish studies have reported the onset of clear intestinal inflammatory responses, including increased expression of pro-inflammatory genes and histological evidence of inflammation^42,43^. However, our qPCR results indicate that this exposure period was still insufficient to induce significant transcriptional changes in the previously discussed genes. Several possible explanations could apply. To start, most studies use soy meal (a rich source of saponins) or soy-derived saponin to induce intestinal inflammation in both larval and adult zebrafish. Here, we used purified saponin derived from *Quillaja saponaria* because it is underexplored in zebrafish. It has been suggested to induce intestinal injury and inflammatory changes in fish, particularly depending on dosage and species^44^. Notably, lower dietary concentrations of *Quillaja* saponins have also been associated with improved growth and feed utilisation in common carp and Nile tilapia^45^, while higher exposures can produce adverse intestinal effects in other fish species^44^. To date, no prominent, direct *in vivo* head-to-head comparison exists between different saponin sources and dosages in adult zebrafish regarding intestinal or overall inflammation, warranting further investigation. More broadly, the host response to bioactive compounds depends not only on source and dosage, but also on exposure duration and route of administration, which influence bioavailability and tissue exposure^46,47^. In this study, the results suggest that the saponin triggered a chemical-induced irritation of the intestinal barrier. As potent surfactants, saponins can irritate the intestinal mucosa by interacting with membrane cholesterol and phospholipids, disrupting membrane integrity, promoting pore formation, and ultimately increasing intestinal permeability^48–50^. Notably, occludin, encoded by *ocnla*, has a regulatory role in tight-junction function and is sensitive to membrane cholesterol and the organisation of cholesterol-rich membrane domains, whereas claudins constitute the principal structural components of tight junctions^51,52^. This could potentially explain the marginal change in *oclna* gene expression and not for *clnd1*. Here, we found that male fish upregulated the expression of both *muc2.1* and *hmgcra* in response to the saponin. The former gene is responsible for mucus secretion^18^, while the latter is involved in *de novo* cholesterol synthesis^19^. The results suggest a compensatory upregulation to offset the saponin-induced chemical trigger. Namely, mucus to coat the lining and mitigate irritation, and 3-hydroxy-3-methylglutaryl-coenzyme A (HMG-CoA) reductase to potentially replenish lost membrane cholesterol.

Here, we show that prior DF intake can sustain intestinal mucosal protection even after cessation. In males, *muc2.1* expression was significantly lower in fish previously fed 15AP than in challenged Controls and was comparable to the unchallenged baseline, suggesting that prior apple pectin intake attenuated the saponin-induced transcriptional effects (**Fig. 10**). Yet we did not observe the same pattern in male fish previously fed the 15C or 30APC diets. We hypothesise that the cellulose may modify the mucus response through its distinct ecological and physical effects within the gut, potentially counteracting or interfering with the effects of AP. In contrast, apple pectin may provide an epigenetic mechanism that may upregulate intestinal mucin production through SCFAs^53^, or independently by promoting gel formation in the gut^54^, which can help neutralise saponin-induced irritation. Additionally, we show that prior DF intake may support the intestinal barrier during saponin-induced stress through different pathways. In males previously fed 15AP or 30APC, *hmgcra* gene expression was significantly lower than in saponin-challenged males (**Fig. 10**). We posit that, even with cellulose added, prior apple pectin intake protected intestinal membrane cholesterol from saponin-induced irritation. Interestingly, our results indicate that female fish appear to be more resilient to saponin than males. This is indicated by consistent gene expression across comparisons and the absence of statistically significant differences following the saponin challenge. Altogether, we show that DFs can leave a sustained imprint on the host even after the dietary intervention ends, potentially contributing to intestinal mucosal protection. However, further work is needed to determine how long these effects persist after DF exposure.

## CONCLUSION

In conclusion, this study highlights that combining DFs can trigger interactions that shape gut microbiome composition in ways not predicted from their isolated effects. Food sources such as fruits, vegetables and legumes often contain a mixture of both soluble and insoluble DFs that can vary drastically in their physicochemical properties. When food is consumed, DFs engage in a dynamic interplay of chemical and physical interactions with other dietary components throughout the digestive tract, driven by properties such as hydration, viscosity and fermentability, thereby modifying the gastrointestinal tract’s physical and chemical environment^55,56^ and shaping the composition and metabolic activity of the gut microbial community^57^. Thus, the health implications of DFs cannot be readily predicted or generalised without accounting for potential interactions with other dietary substrates. Biological responses to DFs also vary across tissues, even within the same host, underscoring the need to better understand the interrelated host-associated factors that shape these responses. Our findings further suggest that DFs may exert a dietary legacy effect, at least at the intestinal barrier level. However, future research should examine additional stressors to determine whether these putative benefits reflect a broader physiological adaptation or remain restricted to the specific challenge. Resolving these dietary and host-dependent interactions will be essential to move beyond broad DF recommendations towards more informed, context-specific dietary strategies.

## METHODS

### Zebrafish husbandry

The zebrafish were sourced from the Zebrafish International Resource Centre (ZIRC) at the University of Oregon, USA. Healthy F2-generation ABWT zebrafish (9 months old) were used in this study. The fish were housed in 3.5 L tanks, with images of gravel underneath each tank, in a recirculating rack system (ZebTEC rack with Active Blue technology, Tecniplast, UK), and kept on a 14:10 hour light:dark cycle. Water temperature was set at 28 °C (range: 26-30 °C), with a pH of 7.25 (range: 7.0-7.5) and a conductivity of 500 μS (range: 300-700 μS). The tank baffles and siphons were washed every other day, and any excess algae were removed throughout the dietary intervention period. We monitored health daily during feeding. Experiments were performed with approval from the UK Home Office under project licence: PP1967239.

### Formulation of diets

We prepared and used two types of dry feed in this study, depending on the experimental phase. For the first phase (dietary intervention), three experimental diets were prepared by supplementing the standard dry feed (Control diet; Zebrafeed 400-600 by SPAROS, Área Empresarial de Marim, Lote C, 8700-221 Olhão, Portugal) with different DFs, as follows: 15% AP (13% w/w), 15% C (13% w/w), and 30% APC (23.1% w/w) (**Table S2**). The same batch of Control diet (25.FO.CE.1102.ZF) was used throughout this study. Fish were fed 5% of their body weight per feed. For 12 fish per tank, the DFs were added to the same amount of Control diet, as follows: (0.3 g Control feed + 0.045 g DF) for the 15% diet and (0.3 g Control feed + 0.045 g DF_1_ + 0.045 g DF_2_) for the 30% diet. The apple pectin (product number 93854) (50 - 75% degree of esterification) and cellulose (product number C6288) were purchased from Sigma-Aldrich (St. Louis, MO, USA). Briefly, the DFs were supplemented into the Control diet by first weighing the desired amount, then adding a small amount of distilled water to create a slurry. This mixture was homogenised and then spread thinly before drying in an oven (50 °C) until bone dry. The next day, the dried sheets were ground in a coffee grinder. The particles were sieved through a 400 μm mesh (Fisher Scientific, Loughborough, Leicestershire, UK). The prepared ground diets were stored in airtight containers in the freezer (−20 °C) until use.

For the second phase of the experiment (saponin challenge), we used Repashy Community Plus (omnivore gel premix) (https://repashy.co.uk/product/community-plus/) as our base food. We incorporated *Quillaja saponaria*-derived saponin (product number: SAE0073-10G) from Sigma into the gel base food at a final concentration of 5 mg/mL (0.5 g saponin powder per 100 mL liquid gel food). Briefly, warm liquid gel food was prepared (3:1 ratio) as per the manufacturer’s instructions before adding the pure saponin powder. Using a magnetic stirrer hotplate (Fischerbrand, UK), the mixture was kept warm (below the boiling point at approx. 50 °C) and well homogenised. Using wide-orifice pipette tips, small gel droplets (approximately 10 µL each) were pipetted onto a non-stick surface on ice. Refer to **Fig. S24** for a photo of the prepared food. The vehicle control is the same base gel food, but without any saponin added. The set gel droplets were kept in the fridge (4-6 °C) until use.

### Experimental design

For the first phase of the experiment (dietary intervention), a total of n = 276 zebrafish (derived from multiple clutches) were randomly assigned to 23 3.5 L tanks after acclimatisation and maintained at a 1:1 sex ratio (six females and six males; n = 12 per tank). The fish in each tank were also randomly assigned to one of the four diets (Control, 15AP, 15C, or 30APC). The weight of 12 fish per tank was measured before the intervention began. The experiment was divided into five blocks, with staggered intervention start dates, each block consisting of four or five tanks (one or two Control tanks and 3 experimental diet tanks). Fish were fed (Monday to Sunday) to satiety (*ad libitum*) with the different diets twice daily (10 AM and 3 PM) at 5% of their body weight each time. The tank water flow was turned off for 10 min to allow undisturbed feeding before switching it back on. Any unconsumed food was allowed to be removed by the zebrafish housing system. All fish received live *Artemia franciscana* enrichment from the Great Salt Lake, Utah, USA (Sep-Art Artemia Cysts, Ocean Nutrition), between the two dry feeds. Each experimental block lasted for three weeks. Refer to **Fig. S1**.

In the second phase (saponin challenge experiment), four fish (two females and two males) from the initial 12 fish were randomly selected to remain in their tanks and advance to the next phase. Similarly, the experiment was divided into five blocks, each consisting of five tanks (one saponin-negative and four saponin-positive). Experimental tanks (saponin-positive) received gel droplets containing saponin. In contrast, the saponin-negative tanks received saponin-free gel food. The tank water flow was turned off for 15 min, after which any leftover gel food was manually syphoned to ensure complete removal within the allocated time. Feeding was also given twice a day (10 AM and 3 PM) from Monday to Sunday, with live *Artemia* enrichment in between. Additionally, in the second phase of the experiment, all tanks were given a concentrated, shell-free *Artemia* solution (decapsulated brine shrimp eggs) purchased from Ocean Aquaculture at approximately 6 PM. Each experimental block lasted for two weeks. Refer to **Fig. S2**.

### Tissue collection and sampling

For the main dietary intervention experiment, we performed dissections once a week (on Mondays) for each block. For dissection, zebrafish were euthanised using a licensed procedure involving hypothermic shock. Briefly, zebrafish were scooped from their tanks and immediately dropped into an ice-cold water bath (< 4 °C), ensuring they did not contact the ice at the bottom. They were left in the ice water bath until the cessation of gill and operculum movement became evident. Blood collection occurred by amputating the caudal tail and aspirating the blood from the dorsal aorta using 75 mm heparinised glass hematocrit tubes (Drummond Scientific Company, PA, USA) before the fish were surgically dissected. From each fish, dissection was performed on ice. Samples of the caudal fin, intestine, gonads, liver, spleen, brain, and blood were harvested without pooling. All tissue and blood samples were immediately flash-frozen in liquid nitrogen and stored at −80 °C until processing.

For the saponin challenge experiment, dissections were performed once a week (on Sundays) for each block. On the dissection days, the shell-free *Artemia* solution feed was skipped, and approximately 3 hrs after their second gel feed, the fish were euthanised by hypothermic shock. Only blood and intestinal samples were collected from each fish for this phase of the study. The intestinal samples were collected directly into tubes containing 600 μL of DNA/RNA shield (Zymo Research, Irvine, CA, USA). Both samples were immediately flash-frozen in liquid nitrogen and stored at −80 °C until processing.

### Quantification of brain, intestinal, and liver metabolites

We followed the same protocol as in our previous study^3^, with some additional optimisation. To enhance tissue lysis, we added approximately 0.6 g of two zirconia/silica bead sizes (0.1 mm and 1.44 mm) to each 1.5 mL conical microtubes with a screw-on O-ring cap (Fisherbrand). The beads and tubes were chilled on dry ice. The zebrafish brain, intestine, and liver (n = 7 per sex per diet per tissue) (refer to **Appendix Fig. S25**), stored at −80 °C, were thawed on ice before weighing on an analytical scale (Fisherbrand) (**Fig. S26**). We then added 1000 µL of 80% ice-cold methanol (80% MeOH + 20% molecular-grade water) to each tube containing the sample and beads. We added twice the methanol (1000 µL instead of 500 µL) to increase the recovery volume after filtration. Samples were homogenised with a FastPrep-24 Classic bead beating grinder and lysis system (MP Biomedicals) using four cycles of 30 sec agitation, each followed by a 90 sec cooling period at −20 °C. The homogenised brain and intestine samples were then incubated at −20 °C for 20 min, then heated to 80 °C for 15 min, and returned to −20 °C for at least 20 min. In contrast, the heating step was omitted for the liver samples, which were kept at −20 °C. Ultimately, all samples were centrifuged (21,000 g) at 4 °C for 5 min. The obtained supernatant was filtered (0.2 µm PTFE filters) into internally threaded cryogenic tubes (Fisherbrand) and stored at −80 °C until analysis.

All samples were concentrated using a Savant SC210A SpeedVac Concentrator (ThermoFisher). Brain and intestinal samples were derivatised in a glass vial by reconstitution with 20 µL HPLC-grade MeOH, followed by the addition of 20 µL of internal standard solution, 10 µL 3-nitrophenylhydrazine (3-NPH) at 50 mM in MeOH with 7% pyridine, and 10 µL 1-Ethyl-3-(3-dimethylaminopropyl) carbodiimide (EDC) at 50 mM in MeOH with 7% pyridine. The solution was vortexed properly and then incubated for 30 min at 37 °C. Finally, the reaction was quenched by adding 20 µL of water with 0.1% formic acid. The column used in all instances was Kinetex® 2.6 µm XB-C18, 50 × 2.1 mm (Phenomenex, Torrance, CA, USA). The SCFAs profile was quantified using a triple-quadrupole LC-MS/MS (Xevo TQ-S Cronos, Waters), controlled by MassLynx v4.1 software for data acquisition, analysis, and management.

Liver samples were reconstituted with 50 µL HPLC-grade MeOH, and 20 µL of internal standard solution (Met-d4 and Trp-d3 at 50 µg/mL in MeOH) was added to the complete extract. Samples were then transferred into glass-vial inserts. Stock solutions of each metabolite (n = 229 across two analytical platforms, with < 5% redundancy between the two metabolite standard stock lists) were used to prepare calibration curves for quantification. Stock solutions were prepared in MeOH and stored at −80 °C, and the internal standard solution was added at a concentration of 50 µg/mL. Samples were analysed using the same Waters Acquity UPLC system and software. For reverse phase analysis, chromatographic separation was performed using a binary gradient of solvent A (Water, 0.1% Formic acid) and solvent B (Acetonitrile, 0.1% Formic acid) at a constant flow rate of 0.3ml/min. A gradient of 0.1% B from 0 to 0.5 min, 10% B from 0.5 to 2.5 min, 15% B at 3.5 min, 35% B at 4.5 min, 45% B at 6.5 min, 55% B at 7 min, and 100% B at 7.5 min. The column was maintained at 100% B until 10 min, followed by a return to 0.1% B at 10.1 min and equilibration under initial conditions until 14 min. Separation was achieved using an ACQUITY Premier HSS T3 column, 2.7 µm (2.1 x 50 mm), maintained at 40 °C. This method was adapted from^58^. For polar compound detection, chromatographic separation (adapted from^59,60^) occurred using a BEH Amide (150 x 2.1 mm, 1.7 µm) using eluent A (10 mM ammonium acetate, 0.1% formic acid) and eluent B (10 mM ammonium acetate, 0.1% formic acid, 90% acetonitrile). At a flow rate of 0.200 mL/min, the LC gradient was as follows: 90% B from to 1.5 min; followed by a linear gradient to 50% B from 1.5 to 11 min; 35% B from 11.5 to 15 min; a return to 90% B at 15.5 min; and equilibration at 90% B from 15.5 to 19 min. The electrospray ionisation (ESI) mode (positive or negative) was selected for each standard based on its ionisation characteristics and optimisation results. Calibration standards were run at the beginning, middle and end of each analytical queue. The analyte: internal standard response ratio was used to create calibration curves and quantify each metabolite.

### RNA extraction and cDNA synthesis

Intestinal samples (n = 10 per sex per treatment) from the saponin experiment were first homogenised by bead beating. Total RNA was extracted using the Quick-RNA MiniPrep Plus (Zymo Research), following the manufacturer’s recommendations. Genomic DNA contamination was removed using the provided DNase I treatment. RNA quality was assessed using NanoDrop™ 8000 Spectrophotometer (Thermo Fisher Scientific) and quantified using the Qubit RNA BR Assay Kit (Invitrogen, Thermo Fisher Scientific, USA). The amount of RNA from intestinal samples was normalised to 1 µg. cDNA synthesis was performed using the UltraScript 2.0 cDNA Synthesis kit (PCR Biosystems, London, UK) as per the manufacturer’s instructions. All cDNA was diluted 1:20 with molecular-grade water and stored at −80 °C until qPCR.

### Gene expression analysis by qPCR

Primers used in this study are presented in **Table S1**. The primer efficiency of each pair was tested using a seven-point standard curve method^61^. The qPCR experiment was performed using PowerUp™ SYBR™ Green Master Mix for qPCR (Applied Biosystems, US) on the QuantStudio™ 7 Flex Real-Time PCR System (384-well; Applied Biosystems, US). The qPCR cycling conditions were as follows: 1 cycle (50°C for 2 min, then 95°C for 2 min) and 40 cycles (95°C for 15 sec, then 60°C for 1 min). This was followed by the conditions needed for the melt curve analysis as defined by the machine. All primers amplified a single product according to melt curve analysis, and amplicon specificity was confirmed by Sanger sequencing (Source Genomics LTD, Cambridge, CB24 9ZR). All 10 µL qPCR reactions (8 µL master mix* and 2 µL cDNA) were performed in duplicate in white 384-well PCR plates. To minimise plate effects, we randomly allocated samples across plates. To correct for run-to-run plate variations, we calculated an inter-run correction for each target gene (*tnfa*, *il1b*, *myd88*, *il6*, *cldn1*, *oclna*, *muc2.1*, *mmp9*, and *hmgcra*). The qPCR results were normalised using the geometric mean of three reference genes: beta-actin (*actb1*), eukaryotic translation elongation factor 1 alpha 1 (*eef1a1*) and ribosomal protein L8 (*rpl8*)^62^. Relative gene expression was first analysed using a linear mixed-effects model, modelling the normalised C_t_ values (ΔC_t_) of the three target genes individually against both fixed and random effects. Then, the comparative (ΔΔCt) method was employed to analyse qPCR data^63^. *(5 µL PowerUp SYBR Green + 0.8 µL 800 nM F. primer + 0.8 µL 800 nM R. primer + 1.4 µL molecular-grade water)

### Genomic DNA extraction, 16S rRNA sequencing, and analysis

Intestinal samples from 10 females and 10 males were selected from each of the four diets (**Fig. S3**). Genomic DNA was extracted using the QIAamp PowerFecal Pro DNA Kit from QIAGEN (Hilden, Germany) as per the manufacturer’s recommendations, which included a bead-beating step using a tissue lyser (MP Biomedicals FastPrep-24 Classic). The extracted genomic DNA was then stored at −20 °C until processing.

16S rRNA amplicon metagenomic sequencing was performed by Novogene (25 Science Park, Milton, Cambridge, CB4 0FW). Once the quality of the sent genomic DNA samples had been validated, the samples were sequenced according to the Novogene workflow. Briefly, amplicon libraries were produced using universal 16S V3-V4 primers (341F: 5’-CCTAYGGGRBGCASCAG-3’ and 806R: 5’-GGACTACNNGGGTATCTAAT-3’), and sequencing was performed using the paired-end 250 base pairs (bp) Illumina NovaSeq 6000 platform with 50 K tags per sample as data output.

The delivered sequencing data had already been processed by Novogene. After demultiplexing, the amplicon sequence data quality was corrected and denoised using DADA2^64^. Taxonomic annotation was performed using the SILVA 138.1 database^65^. Bioinformatics analysis was conducted using a combination of QIIME2^66^ and *phyloseq*^67^ in RStudio. α-diversity metrics, namely Observed, FaithPD, Shannon, and Simpson, were used to assess within-group diversity. We assessed the effects of “Diet”, “Sex”, and their interaction on unweighted and weighted UniFrac β-diversity and Aitchison distance β-diversity using PERMANOVA (9999 permutations). For Aitchison distance β-diversity, we filtered unrarefied raw reads to exclude low-prevalence and low-abundance taxa, retaining features detected in ≥10% of samples (prevalence) and with a mean relative abundance ≥0.005%. We then applied a centred log ratio (CLR) transformation with a pseudocount of 1, controlling for differences in library size and suitable for the compositional nature of microbiome data. For the component analysis, diet was encoded as a 2 × 2 factorial design based on the presence or absence of apple pectin (AP) and cellulose (C): Control = AP absent/C absent; 15AP = AP present/C absent; 15C = AP absent/C present; and 30APC = AP present/C present. We performed differential abundance analysis using ANCOM-BC2 (models: Diet + Sex + Block), with a minimum library size of 100, a prevalence threshold of 0.1, and Benjamini-Hochberg (BH) for multiple comparison adjustment. Predicted functional profile analysis was performed using PICRUSt2^68^. Individual predicted pathways were analysed using limma^69^ on CLR-transformed abundances, with “Diet” and “Sex” accounted for in the models. For the microbiome-metabolomics correlation analysis, we selected MetaCyc (https://biocyc.org/meta/) pathways with established primary relevance to acetate, propionate, and butyrate metabolism from the predicted pathway profiles. Pathway abundances were converted to relative abundance within each sample and summarised by Diet × Sex. Acetate-associated pathways included: P124-PWY (Bifidobacterium shunt), P461-PWY (hexitol fermentation to lactate, formate, ethanol and acetate), P163-PWY (L-lysine fermentation to acetate and butanoate) and PWY-5100 (pyruvate fermentation to acetate and lactate II); propionate-associated pathways included: P108-PWY (pyruvate fermentation to propanoate I), PWY-5088 (L-glutamate degradation VIII to propanoate) and PWY-7013 (L-1,2-propanediol degradation); and butyrate-associated pathways included: P163-PWY (L-lysine fermentation to acetate and butanoate), PWY-5676 (acetyl-CoA fermentation to butanoate II) and CENTFERM-PWY (pyruvate fermentation to butanoate)^70^. Because the propionate-associated pathways showed broadly coherent responses, we standardised their group-level abundances as z-scores and averaged them to generate a composite propionate pathway score. In contrast, we analysed acetate- and butyrate-associated pathways separately because their responses were pathway-specific. Specifically, for propionate, we retained P108-PWY as the primary pathway-specific marker. We also generated a composite predicted propionate pathway score from P108-PWY, PWY-5088, and PWY-7013. Additionally, we grouped (P461-PWY and P124-PWY) and (P163-PWY and PWY-5100) for the acetate analysis because they behaved similarly, but we kept the three butyrate-associated pathways separate.

### Growth metrics and survival parameters

At the start and end of the three-week dietary intervention, fish growth metrics and survival parameters were assessed (refer to **Fig. S27**). The mean weight per fish in each tank was divided by the total number of fish as follows:

- Initial mean weight per fish in tank = (Initial tank weight)/12 fish per tank
- Final mean weight per fish in tank = (Final tank weight)/ fish survivors

The following formula for the growth metrics was used:

i. Weight difference = (final mean weight) - (initial mean weight)
ii. % weight gain = (weight difference)/ (initial mean weight) X 100
iii. Specific growth rate (SGR) = ([(ln_final mean weight_)-(ln_initial mean weight_)]/ days) X 100

where ln = natural log and days = 21

Survival for each tank was analysed as count data as follows:

- Fish survivors: alive at 3 weeks
- Fish dead: 12 - fish survivors

### Statistical analysis

Statistical analyses were performed using R (version 4.6.1)^71^. Unless otherwise stated, descriptive data are presented as mean ± standard error of the mean (SEM). For lmer, degrees of freedom were estimated using the Satterthwaite approximation. Model assumptions were evaluated using appropriate diagnostic procedures. Statistical tests were two-sided, and significance was defined as *p* < 0.05. Intestinal and brain SCFAs were analysed together in a full model. Concentrations were initially log10-transformed, and linear mixed-effects models were fitted with “Diet”, “Sex”, “Tissue” and their interactions as fixed effects. We applied a Box-Cox transformation to valerate (λ ≈ −0.36), butyrate (λ ≈ 0.49), isobutyrate (λ ≈ 0.42), and caproate (λ ≈ 0.47) to meet linear mixed model assumptions. Where applicable, multiple comparisons were performed and *p* values were adjusted for multiple testing. Holm’s method was applied to families of planned contrasts, Tukey’s method to complete pairwise comparisons among diet levels, and the Benjamini-Hochberg false discovery rate (FDR) procedure to metabolomics analyses involving multiple metabolites. Adjusted *p* values are reported throughout as *p*_adj_, with multiple comparisons performed as specified.

