## Supplementary Material for "Dietary fibre combinations modulate gut microbiome composition and metabolic responses in zebrafish, with lasting effects on intestinal barrier gene expression"

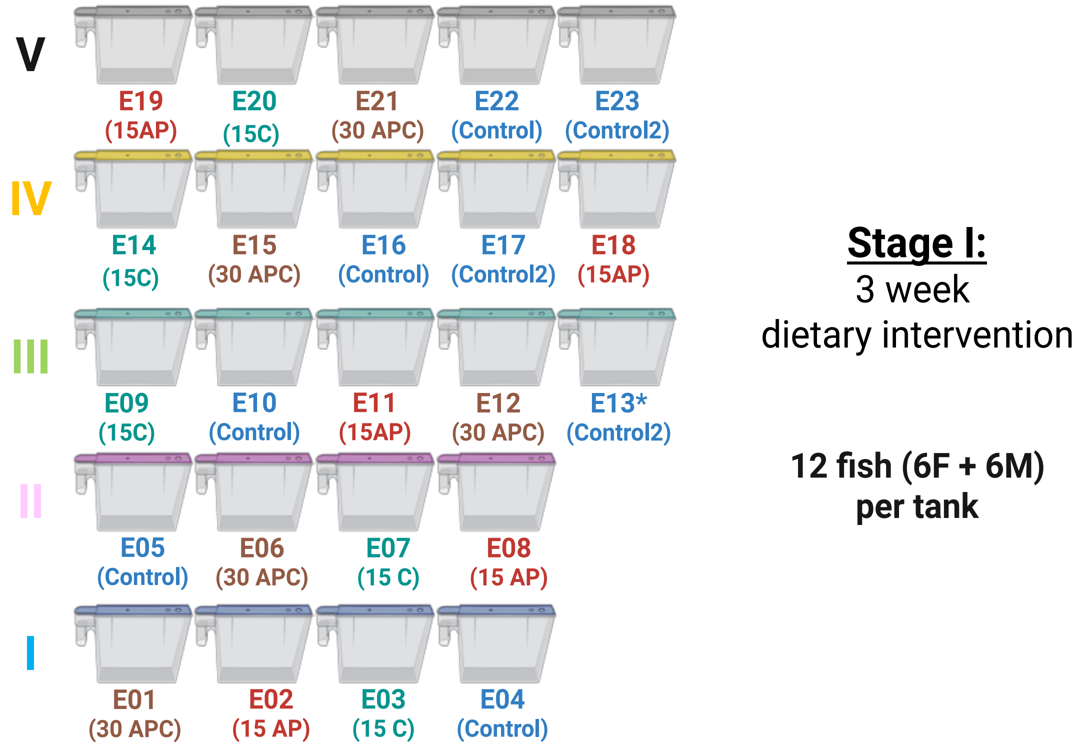


**Fig. S1:** Experimental design setup for the dietary intervention.

Female and male zebrafish were housed 1:1 for a total of 12 fish per tank. Roman numerals denote the block number. Each tank was assigned an ID and a diet. *Tank E13 was used to top up tanks E04 and E05 for the saponin experiment. Tanks E04, E05, E10, E16, and E22 were the Control diet groups for the first stage of the study. For the saponin experiment, fish from the two Control tanks (blocks III, IV, and V) were randomly shuffled to generate Control_Sap-positive and Control_Saponin-negative tanks per block. Created in **https://BioRender.com**

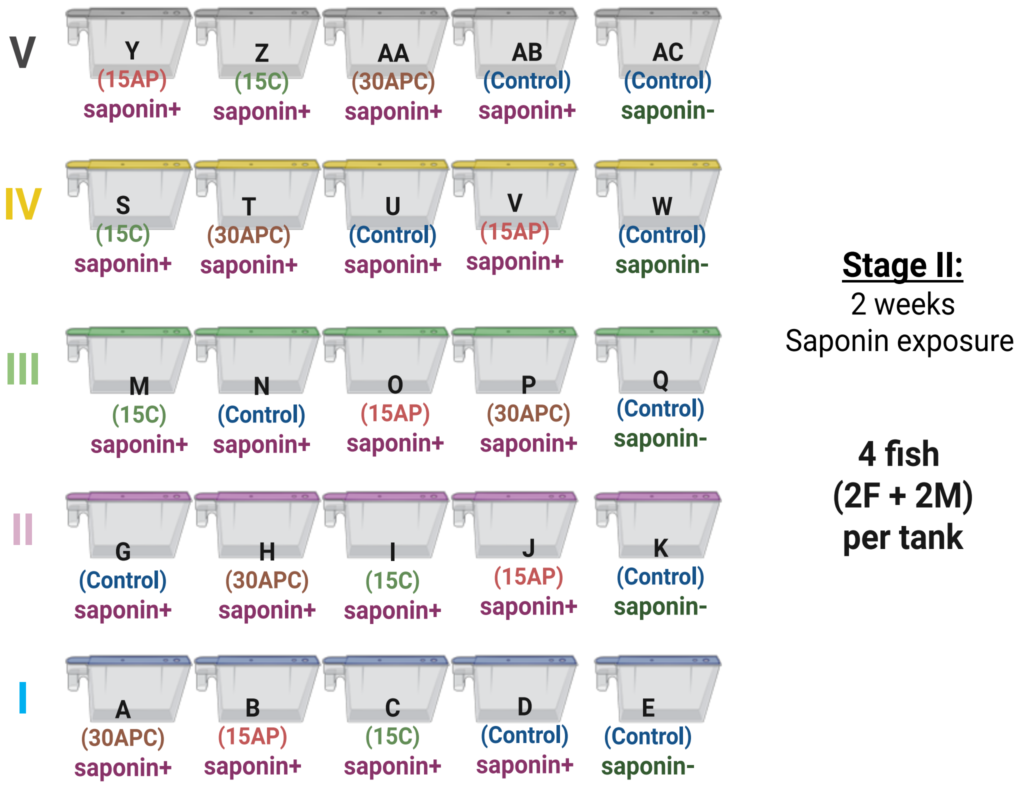


**Fig. S2:** Experimental design setup for the saponin challenge.

Female and male zebrafish were housed 1:1 for a total of 4 fish per tank. Roman numerals denote the block number. Each tank was assigned an ID and a treatment. Created in **https://BioRender.com**

| Diet | Sex | n |
|---|---|---|
| Control | F | 10 |
| Control | M | 10 |
| 15AP | F | 10 |
| 15AP | M | 10 |
| 15C | F | 9 |
| 15C | M | 9 |
| 30APC | F | 10 |
| 30APC | M | 10 |

**Fig. S3:** Intestinal sample size for 16S rRNA sequencing.

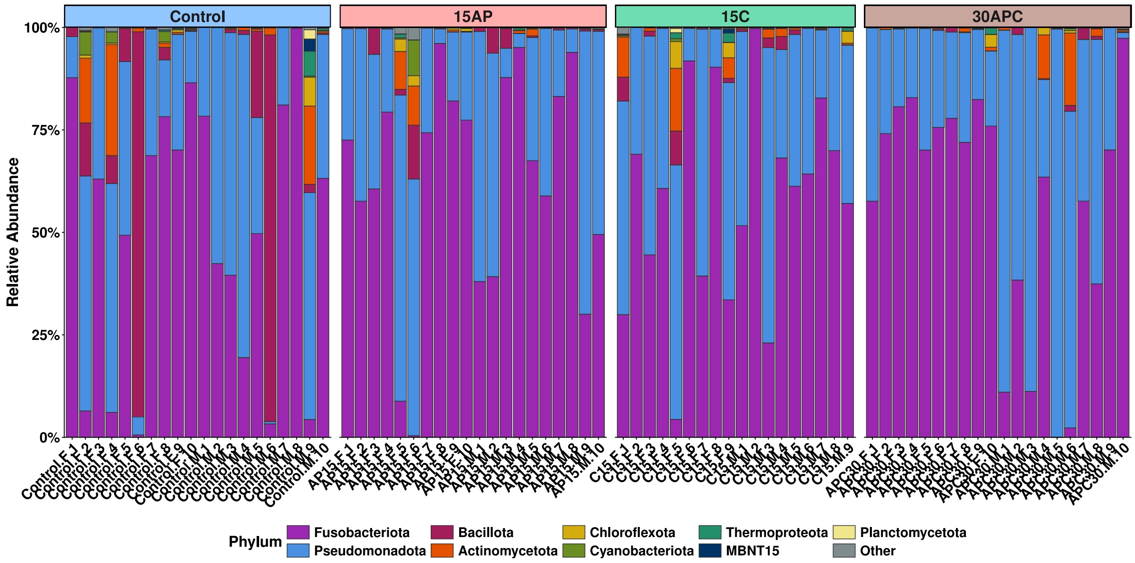


**Fig. S4:** Relative abundance of bacterial phyla.

Fusobacteriota and Pseudomonadota (Proteobacteria) were the two most abundant phyla across samples

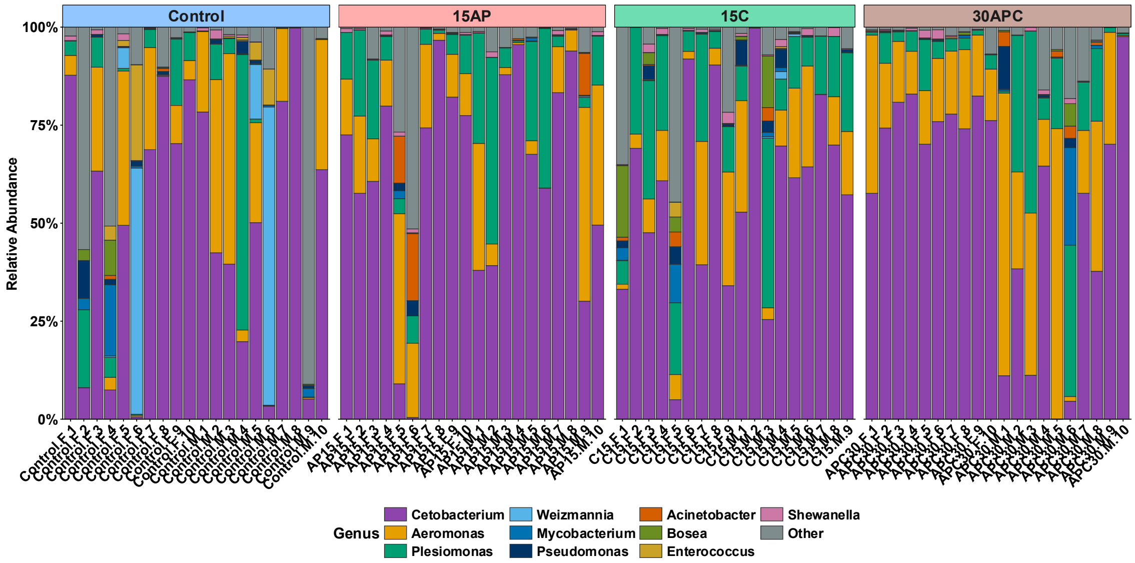


**Fig. S5:** Relative abundance at the Genus level.

*Cetobacterium*, *Aeromonas*, and *Plesiomonas* were the core genera across the samples.

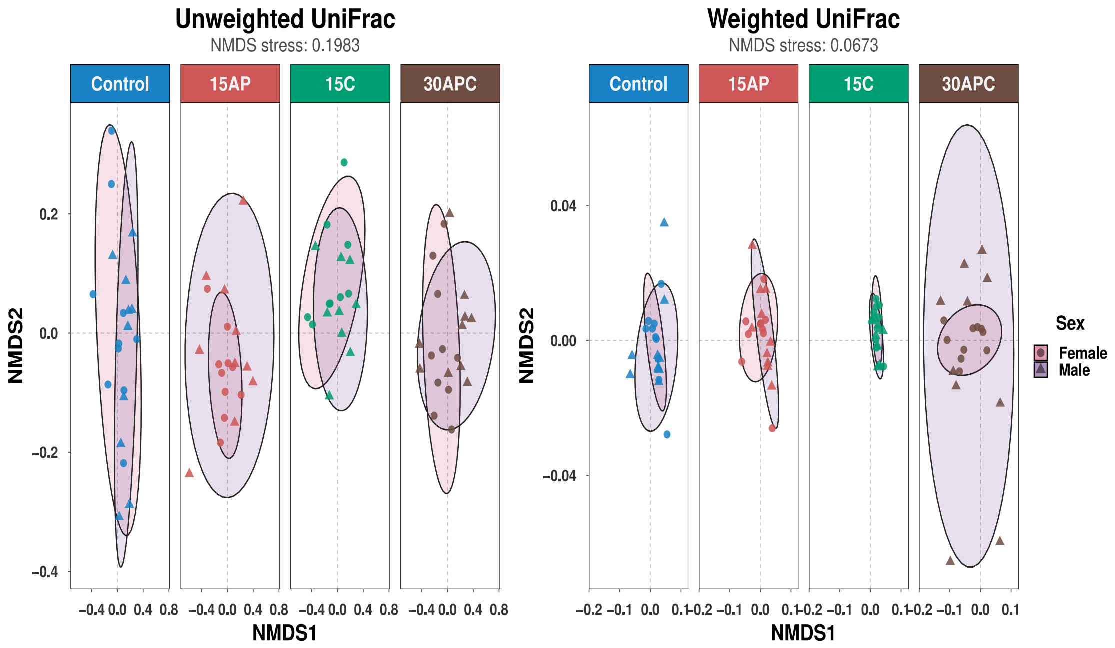


**Fig. S6:** Unweighted and weighted UniFrac β-diversity grouped by sex and faceted by diet. Non-metric multidimensional scaling (NMDS) plots showing variation in gut microbial community composition between females and males within each dietary group. Stress values indicate the goodness of fit.

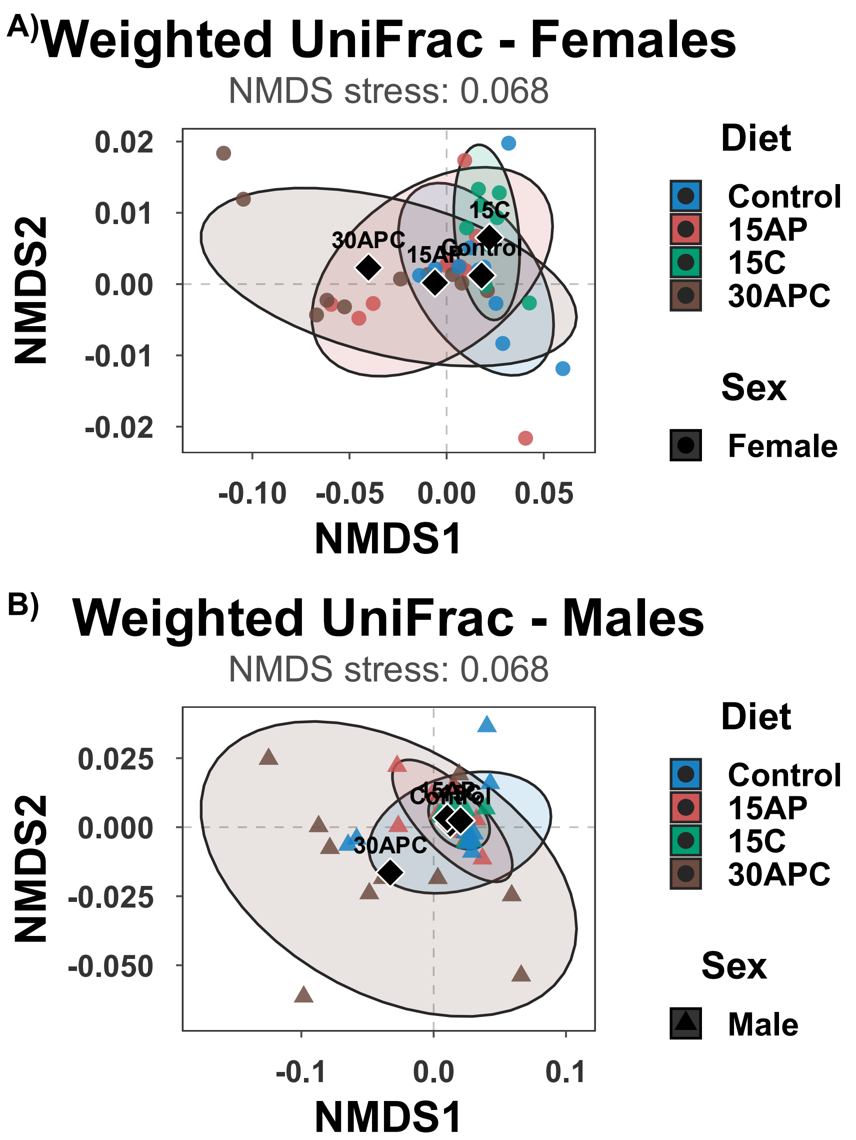


**Fig. S7: Weighted UniFrac β-diversity by diet, stratified by sex.**

(**A**) Weighted UniFrac β-diversity in females. (**B**) Weighted UniFrac β-diversity in males.

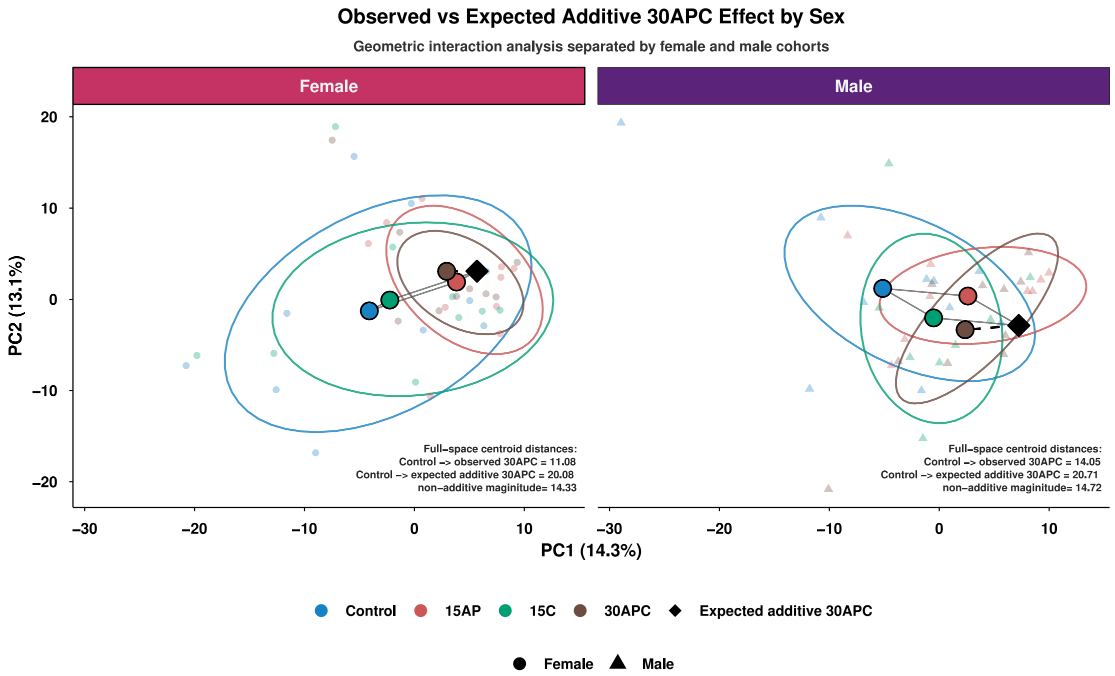


**Fig. S8:** Observed versus expected additive effects of the 30APC diet on gut microbiome composition, stratified by sex.

Aitchison distances quantified compositional differences between the observed 30APC microbiome and the expected composition under an additive response.

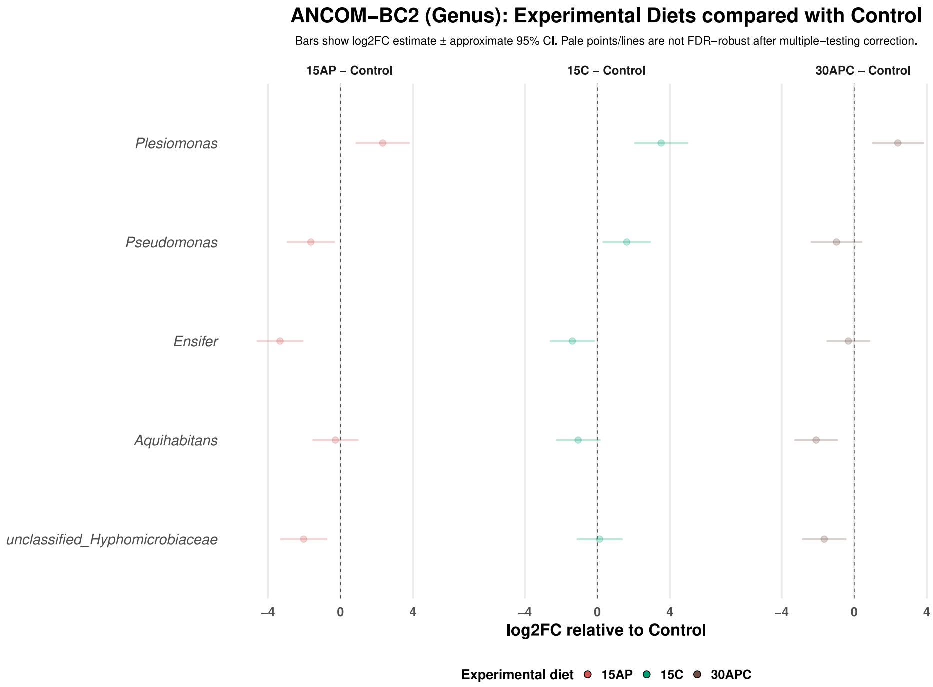


**Fig. S9:** Differential abundance analysis comparing experimental to Control.

Each point represents the model-estimated log2FC, and the horizontal error bar represents its 95% confidence interval

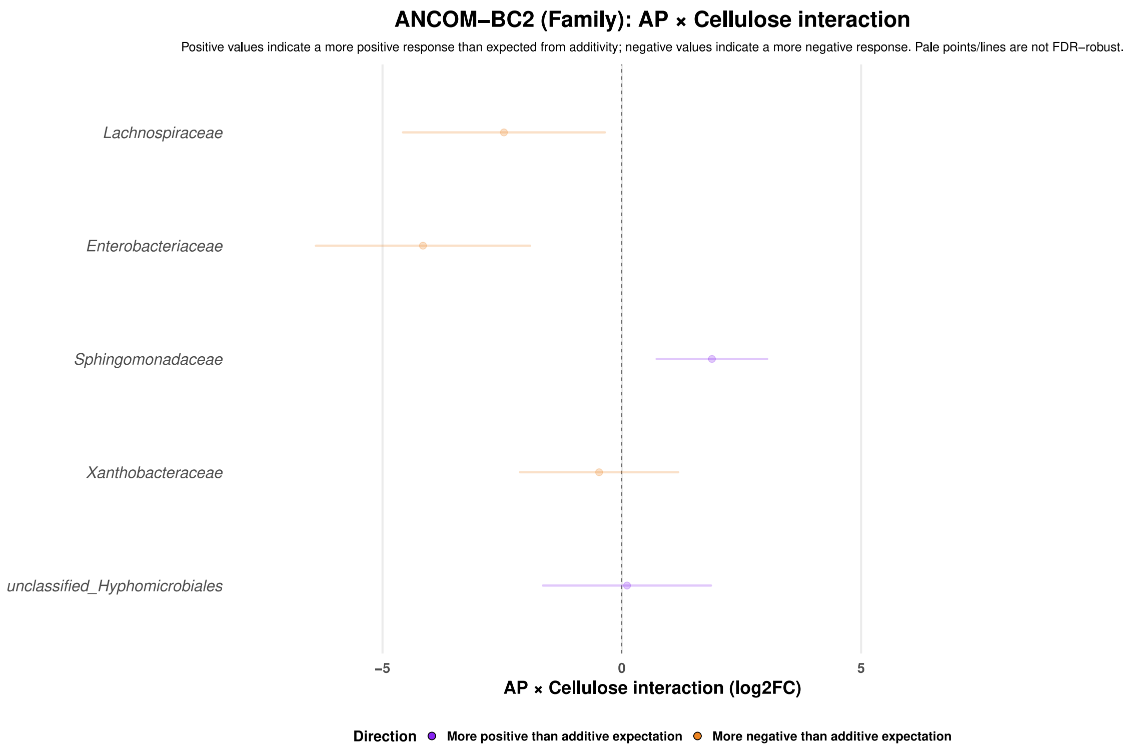


**Fig. S10:** Differential abundance analysis of the AP × C interaction at the family level.

Positive values indicate higher abundance than expected under additivity; negative values indicate lower abundance than expected under additivity. Each point represents the model-estimated log2FC, and the horizontal error bar represents its 95% confidence interval. No individual family deviated significantly from the additive expectation after FDR correction.

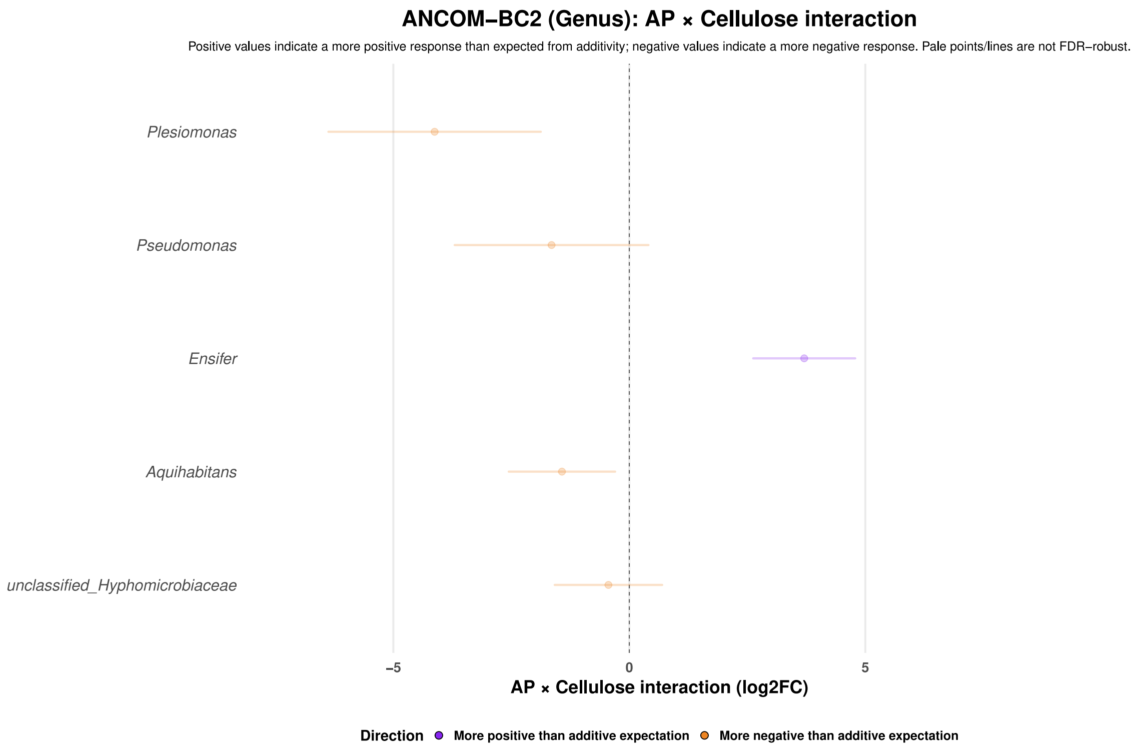


**Fig. S11:** Differential abundance analysis of the AP × C interaction at the genus level.

Positive values indicate higher abundance than expected under additivity; negative values indicate lower abundance than expected under additivity. Each point represents the model-estimated log2FC, and the horizontal error bar represents its 95% confidence interval. No individual genera deviated significantly from the additive expectation after FDR correction.

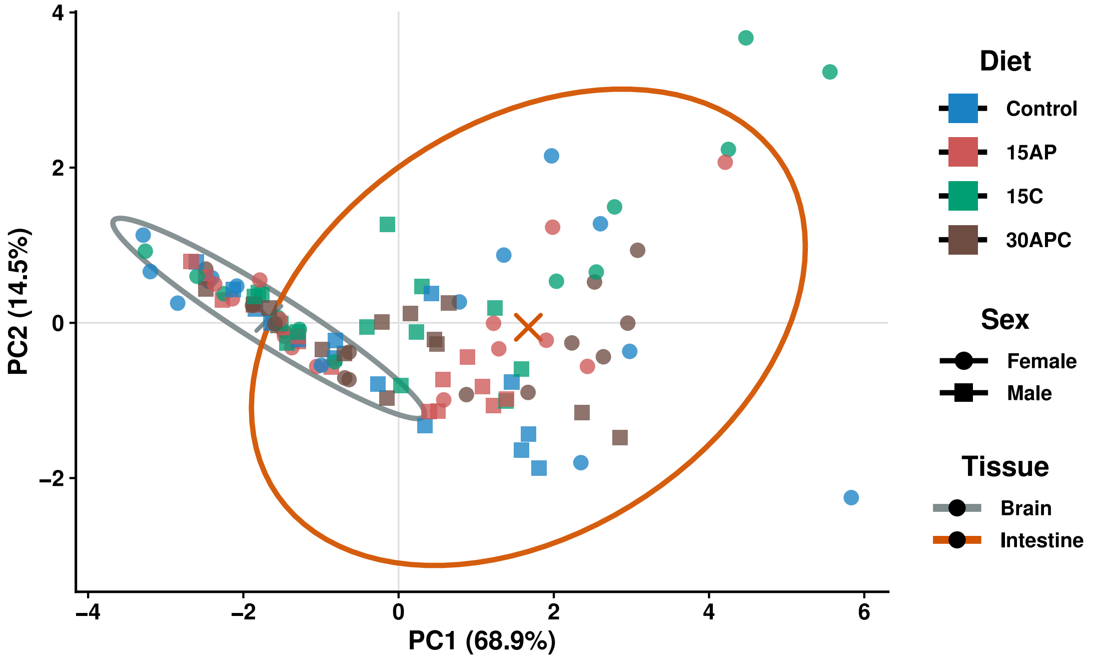


**Fig. S12:** PCA of quantified SCFAs showing clustering by tissue.

### Combined SCFAs — final linear mixed-effects models

Type III F-tests with Satterthwaite denominator degrees of freedom

| Outcome | Term | Num df | Den df | *F* | *p* |
|---|---|---|---|---|---|
| Acetate | Diet | 3 | 42.55 | 0.78 | 0.511 |
| | Sex | 1 | 44.37 | 2.22 | 0.143 |
| | Tissue | 1 | 47.14 | 186.58 | <0.001 |
| | Diet × Sex | 3 | 43.32 | 5.63 | 0.002 |
| | Diet × Tissue | 3 | 46.97 | 3.24 | 0.030 |
| | Sex × Tissue | 1 | 47.14 | 22.64 | <0.001 |
| | Diet × Sex × Tissue | 3 | 46.97 | 0.19 | 0.900 |
| Butyrate | Diet | 3 | 43.03 | 1.95 | 0.136 |
| | Sex | 1 | 45.00 | 2.71 | 0.107 |
| | Tissue | 1 | 47.18 | 247.74 | <0.001 |
| | Diet × Sex | 3 | 43.87 | 2.22 | 0.099 |
| | Diet × Tissue | 3 | 47.03 | 0.42 | 0.740 |
| | Sex × Tissue | 1 | 47.18 | 9.96 | 0.003 |
| | Diet × Sex × Tissue | 3 | 47.03 | 1.61 | 0.199 |
| Caproate | Diet | 3 | 89.90 | 0.57 | 0.635 |
| | Sex | 1 | 92.97 | 8.52 | 0.004 |
| | Tissue | 1 | 88.51 | 286.72 | <0.001 |
| | Diet × Sex | 3 | 90.96 | 1.12 | 0.346 |
| | Diet × Tissue | 3 | 88.39 | 2.74 | 0.048 |
| | Sex × Tissue | 1 | 88.51 | 12.80 | <0.001 |
| | Diet × Sex × Tissue | 3 | 88.39 | 3.09 | 0.031 |
| Isobutyrate | Diet | 3 | 90.52 | 2.70 | 0.050 |
| | Sex | 1 | 93.47 | 1.59 | 0.210 |
| | Tissue | 1 | 88.96 | 217.19 | <0.001 |
| | Diet × Sex | 3 | 91.50 | 2.42 | 0.071 |
| | Diet × Tissue | 3 | 88.81 | 0.08 | 0.968 |
| | Sex × Tissue | 1 | 88.96 | 7.67 | 0.007 |
| | Diet × Sex × Tissue | 3 | 88.81 | 0.90 | 0.446 |
| Propionate | Diet | 3 | 40.85 | 1.00 | 0.404 |
| | Sex | 1 | 41.68 | 20.24 | <0.001 |
| | Tissue | 1 | 47.40 | 618.64 | <0.001 |
| | Diet × Sex | 3 | 41.23 | 1.97 | 0.133 |
| | Diet × Tissue | 3 | 47.18 | 0.45 | 0.721 |
| | Sex × Tissue | 1 | 47.40 | 49.09 | <0.001 |
| | Diet × Sex × Tissue | 3 | 47.18 | 4.29 | 0.009 |
| Valerate | Diet | 3 | 15.93 | 2.64 | 0.085 |
| | Sex | 1 | 84.55 | 18.46 | <0.001 |
| | Tissue | 1 | 79.50 | 16.02 | <0.001 |
| | Diet × Sex | 3 | 84.44 | 0.84 | 0.478 |
| | Diet × Tissue | 3 | 79.50 | 4.80 | 0.004 |
| | Sex × Tissue | 1 | 79.50 | 18.54 | <0.001 |
| | Diet × Sex × Tissue | 3 | 79.50 | 1.09 | 0.357 |

**Fig. S13:** Type III ANOVA output for combined-tissue SCFA linear mixed-effects models.

### SCFA component analysis — final linear mixed-effects models

Selected Type III F-tests with Satterthwaite denominator degrees of freedom

| Outcome | Term | Num df | Den df | *F* | *p* |
|---|---|---|---|---|---|
| Acetate | Apple pectin component | 1 | 42.56 | 0.08 | 0.776 |
| | Cellulose component | 1 | 42.15 | 0.79 | 0.380 |
| | Sex | 1 | 43.04 | 7.77 | 0.008 |
| | Tissue | 1 | 46.82 | 85.11 | <0.001 |
| | Apple pectin component × Cellulose component | 1 | 42.23 | 0.72 | 0.402 |
| Butyrate | Apple pectin component | 1 | 43.03 | 3.07 | 0.087 |
| | Cellulose component | 1 | 42.59 | 2.76 | 0.104 |
| | Sex | 1 | 43.65 | 1.67 | 0.204 |
| | Tissue | 1 | 46.90 | 69.19 | <0.001 |
| | Apple pectin component × Cellulose component | 1 | 42.64 | 0.65 | 0.426 |
| Caproate | Apple pectin component | 1 | 89.96 | 1.19 | 0.279 |
| | Cellulose component | 1 | 89.10 | 0.01 | 0.922 |
| | Sex | 1 | 90.43 | 0.03 | 0.858 |
| | Tissue | 1 | 88.31 | 106.22 | <0.001 |
| | Apple pectin component × Cellulose component | 1 | 89.22 | 0.17 | 0.677 |
| Isobutyrate | Apple pectin component | 1 | 90.56 | 1.39 | 0.242 |
| | Cellulose component | 1 | 89.70 | 2.96 | 0.089 |
| | Sex | 1 | 91.25 | 2.97 | 0.088 |
| | Tissue | 1 | 88.72 | 55.69 | <0.001 |
| | Apple pectin component × Cellulose component | 1 | 89.79 | 0.01 | 0.940 |
| Propionate | Apple pectin component | 1 | 39.94 | 0.59 | 0.448 |
| | Cellulose component | 1 | 40.57 | 0.96 | 0.332 |
| | Sex | 1 | 40.86 | 0.03 | 0.868 |
| | Tissue | 1 | 46.97 | 164.74 | <0.001 |
| | Apple pectin component × Cellulose component | 1 | 40.10 | 0.00 | 0.989 |
| Valerate | Apple pectin component | 1 | 15.72 | 0.06 | 0.809 |
| | Cellulose component | 1 | 16.15 | 6.47 | 0.022 |
| | Sex | 1 | 85.69 | 3.41 | 0.068 |
| | Tissue | 1 | 79.02 | 0.01 | 0.904 |
| | Apple pectin component × Cellulose component | 1 | 15.94 | 2.44 | 0.138 |

**Fig. S14:** Type III ANOVA output from SCFA component-analysis linear mixed-effects models.

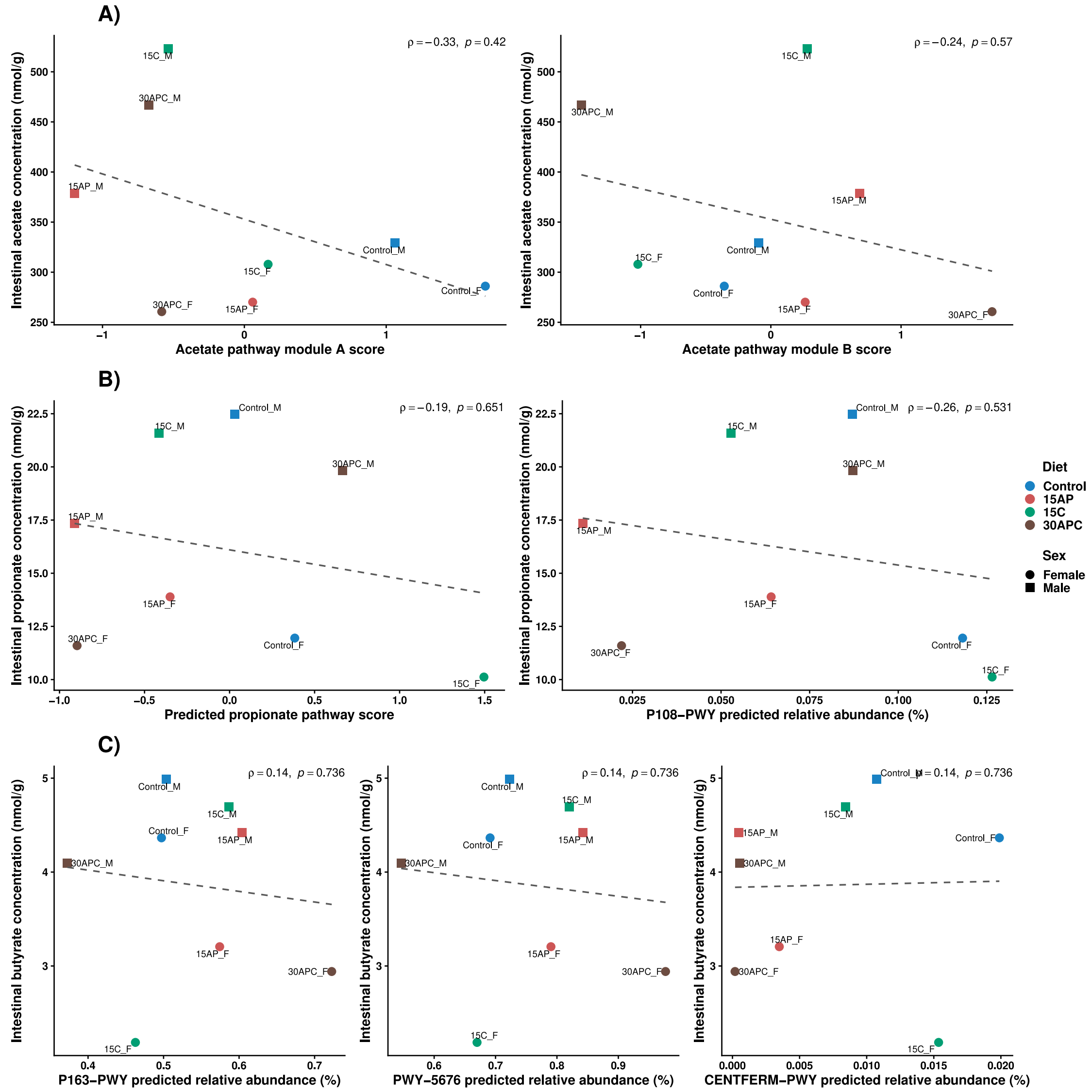


**Fig. S15:** Group-level Spearman correlations between intestinal concentrations of acetate, propionate, and butyrate and the selected microbial predicted pathways.

(**A**) Correlations between intestinal acetate and predicted microbial pathways. (**B**) Correlations between intestinal propionate and predicted microbial pathways. (**C**) Correlations between intestinal butyrate and predicted microbial pathways. Correlations were generally weak (ρ) and not statistically significant ($p > 0.05$).

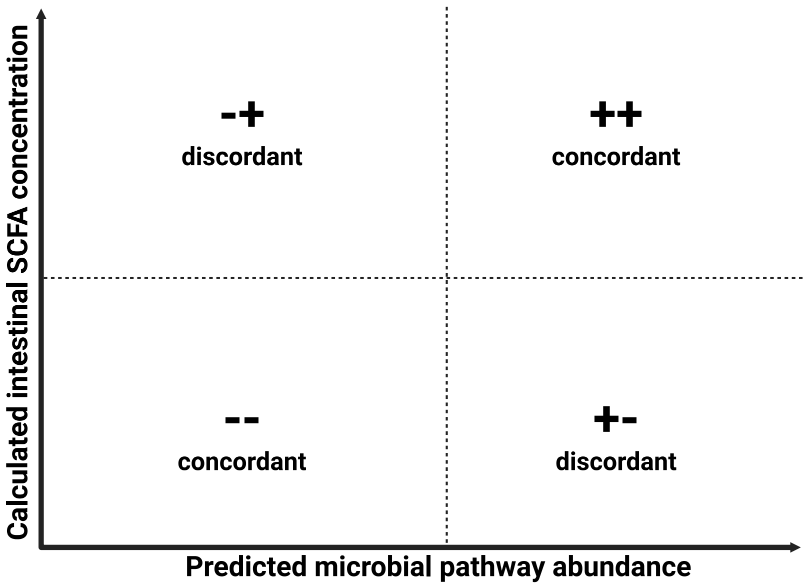


**Fig. S16:** Four-quadrant plot map.

Four-quadrant plot, where the upper-right quadrant indicates a directionally concordant increase and the lower-left quadrant a directionally concordant decrease.

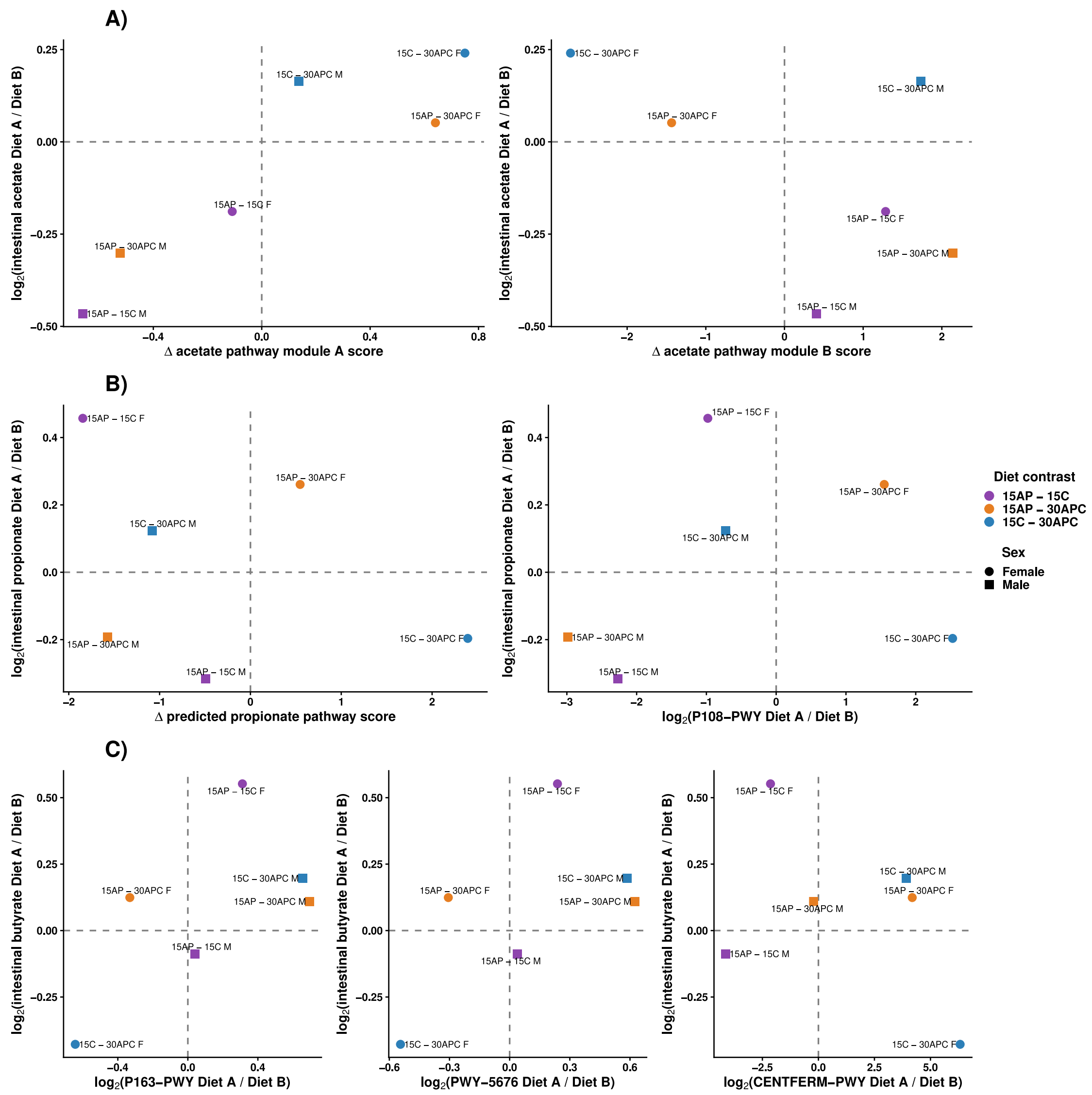


**Fig. S17:** Direction-concordance analysis comparing the pairwise experimental diets in the intestine.

Direction concordance indicates whether predicted microbial functions changed in the same or opposing directions across experimental diet contrasts. (**A**) Predicted microbial functional response associated with intestinal acetate. (**B**) Predicted microbial functional response associated with intestinal propionate. (**C**) Predicted microbial functional response associated with intestinal butyrate. For each comparison, Diet A is the numerator, and Diet B is the denominator in the log2(Diet A/Diet B) ratio. Positive values indicate a higher response in Diet A relative to Diet B, whereas negative values indicate a higher response in Diet B relative to Diet A.

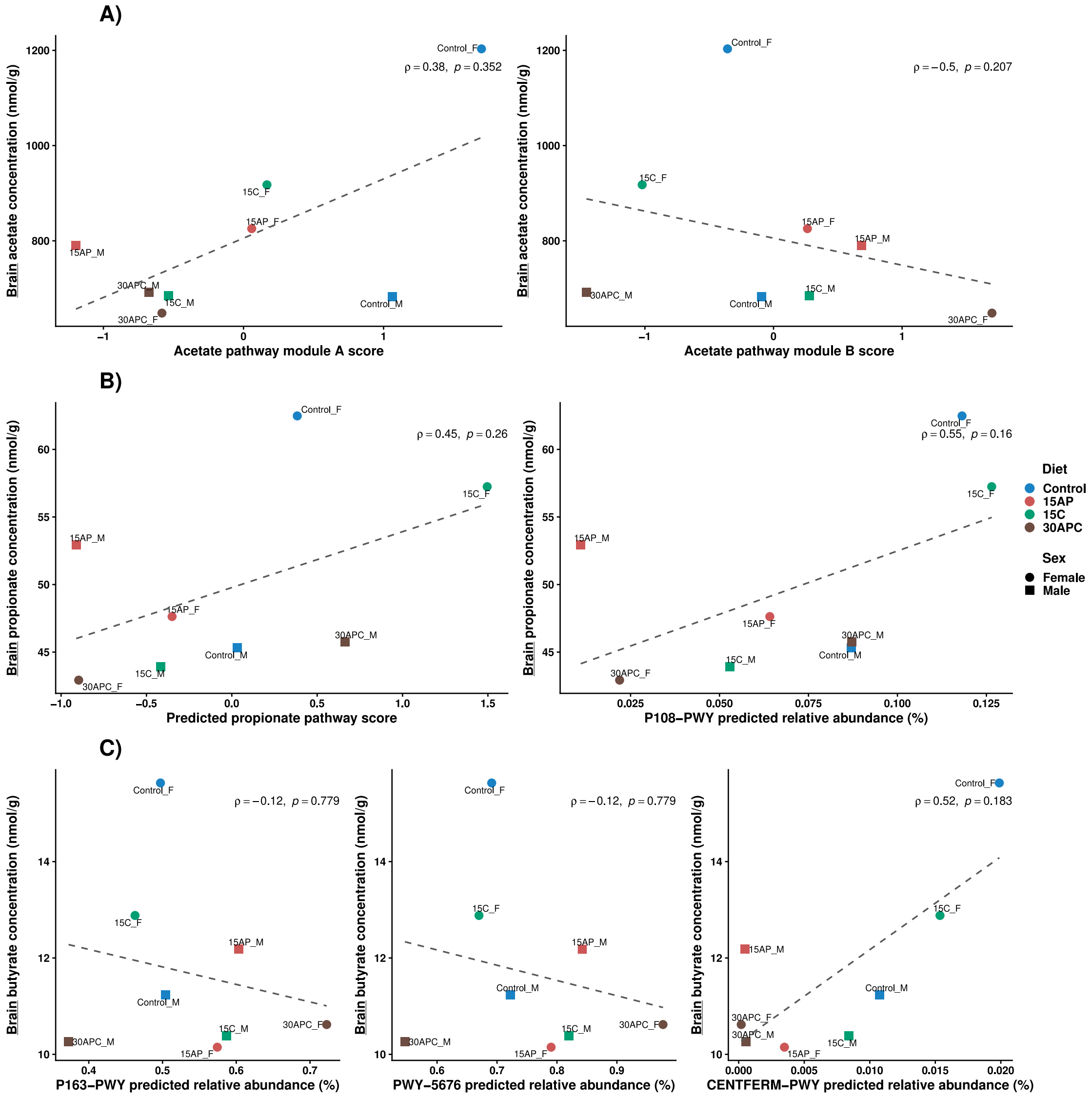


**Fig. S18:** Group-level Spearman correlations between brain concentrations of acetate, propionate, and butyrate and the selected microbial predicted pathways.

(**A**) Correlations between brain acetate and predicted microbial pathways. (**B**) Correlations between brain propionate and predicted microbial pathways. (**C**) Correlations between brain butyrate and predicted microbial pathways.

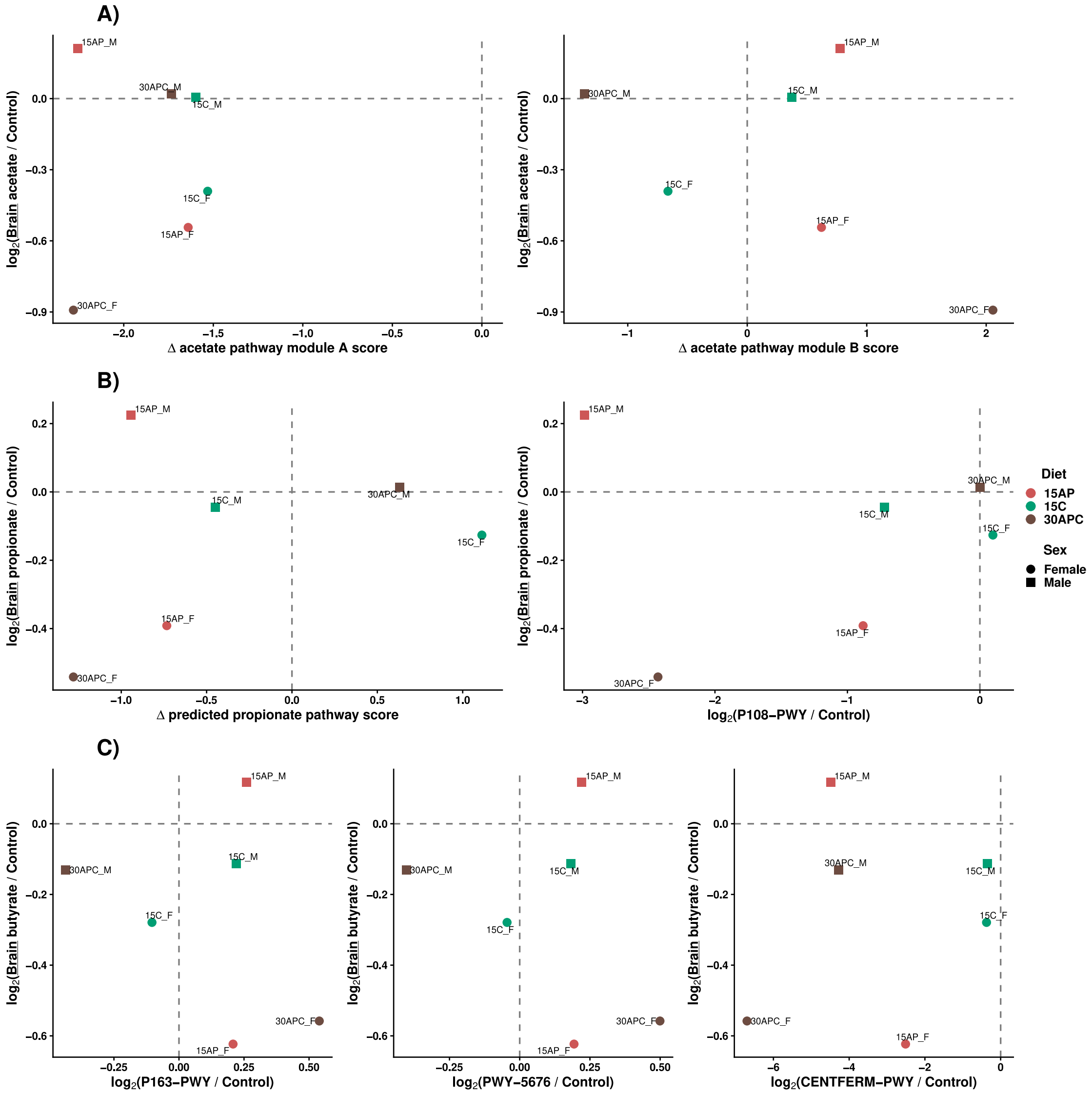


**Fig. S19:** Direction-concordance analysis comparing the experimental diets relative to Control in the brain.

(**A**) Predicted microbial functional response associated with brain acetate. (**B**) Predicted microbial functional response associated with brain propionate. (**C**) Predicted microbial functional response associated with brain butyrate. For each contrast, the response direction was defined relative to Control, with positive values indicating an increase and negative values indicating a decrease relative to Control.

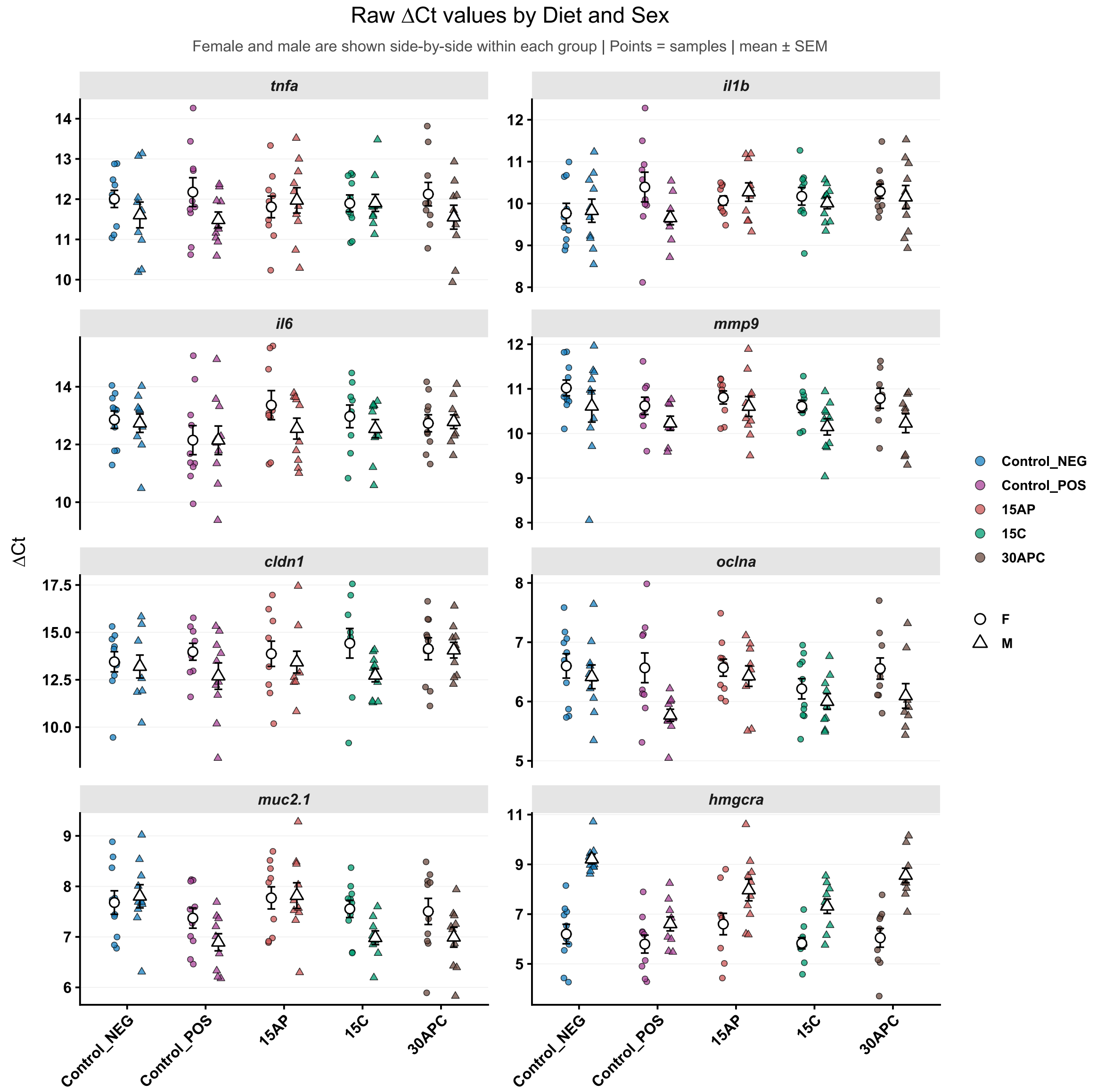


**Fig. S20:** Normalised intestinal gene expression of target genes.

Blood raw ΔCt values of the target genes (*tnfa*, *il1b*, *il6, mmp9, cldn1, ocnla, muc2.1,* and *hmgcra*) normalised to the three reference genes (*actb1*, *eef1a1*, and *rpl8)*. The centre dots and error bars indicate the mean ± standard error of the mean (SEM). F denotes females, and M denotes males.

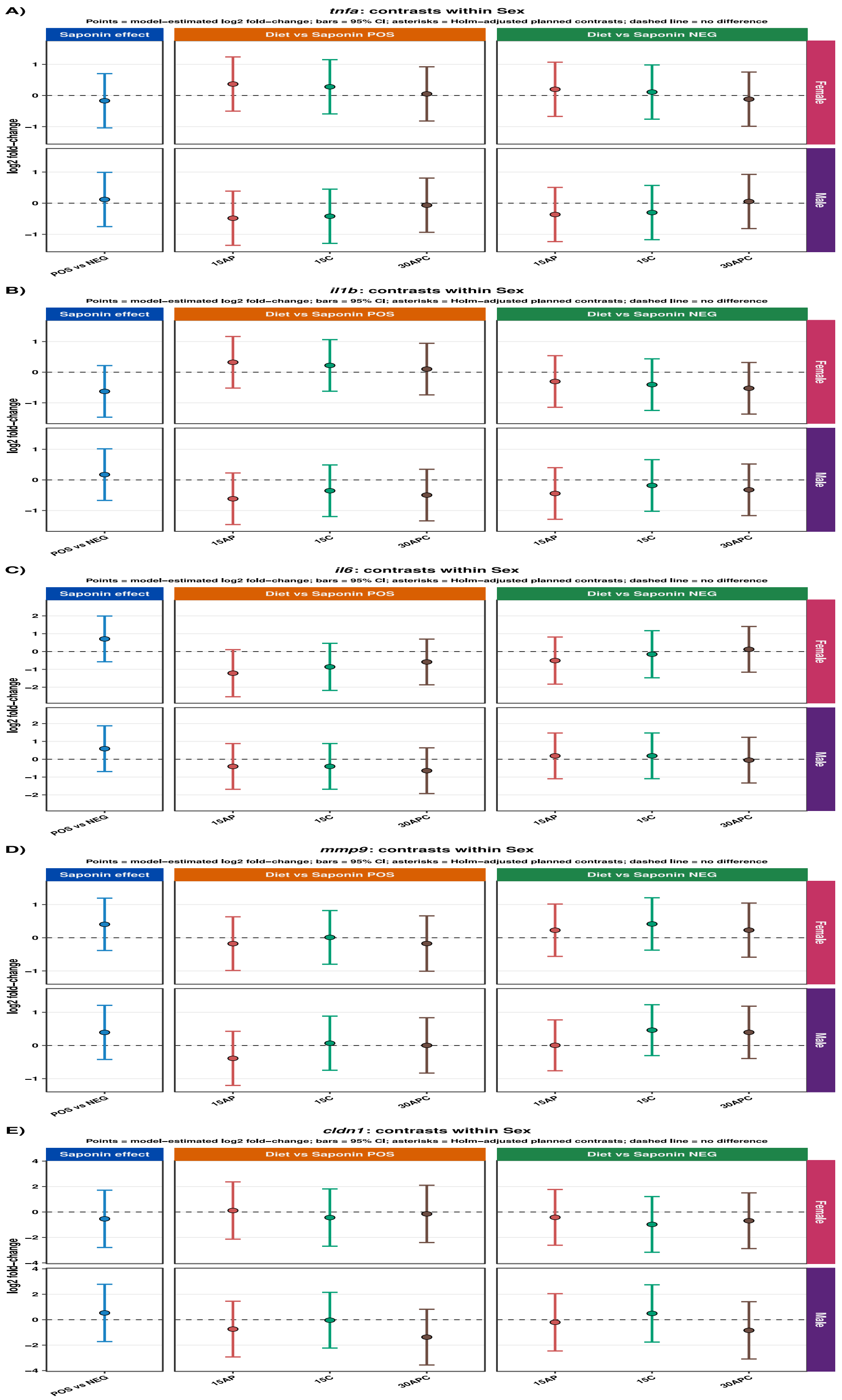


**Fig. S21:** Intestinal gene expression of *tnfa*, *il1b*, *il6*, *mmp9,* and *cldn1* under the various contrasts, stratified by sex.

(**A**) Intestinal gene expression patterns of *tnfa*. **(B)** Intestinal gene expression patterns of *il1b*. **(C)** Intestinal gene expression patterns of *il6*. **(D)** Intestinal gene expression patterns of *mmp9.* **(E)** Intestinal gene expression patterns of *cldn1*. Each point is the model-estimated log2FC, and the vertical error bar is its 95% confidence interval.

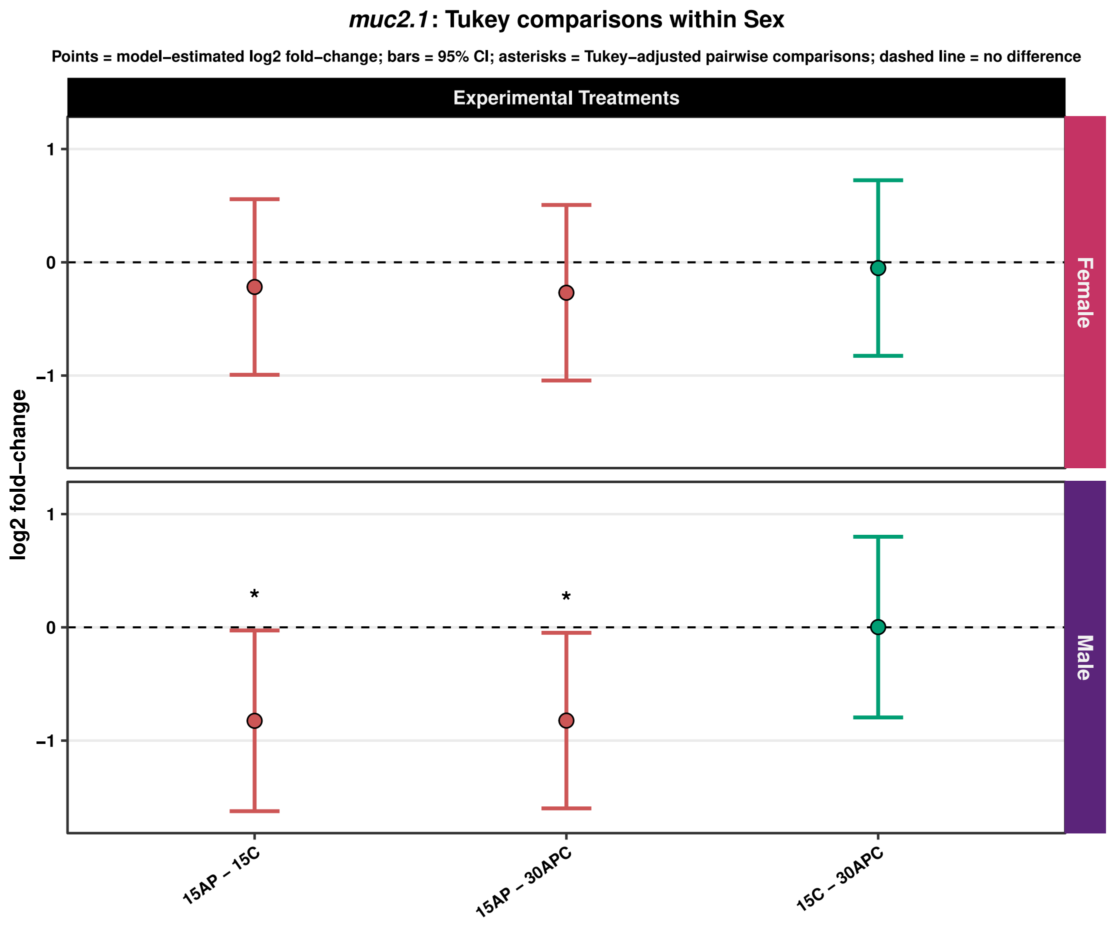


**Fig. S22:** Intestinal *muc2.1* gene expression across experimental diet comparisons and faceted by sex.

Diet pairwise contrasts of *muc2.1* expression. Each point is the model-estimated log2FC, and the vertical error bar is its 95% confidence interval.

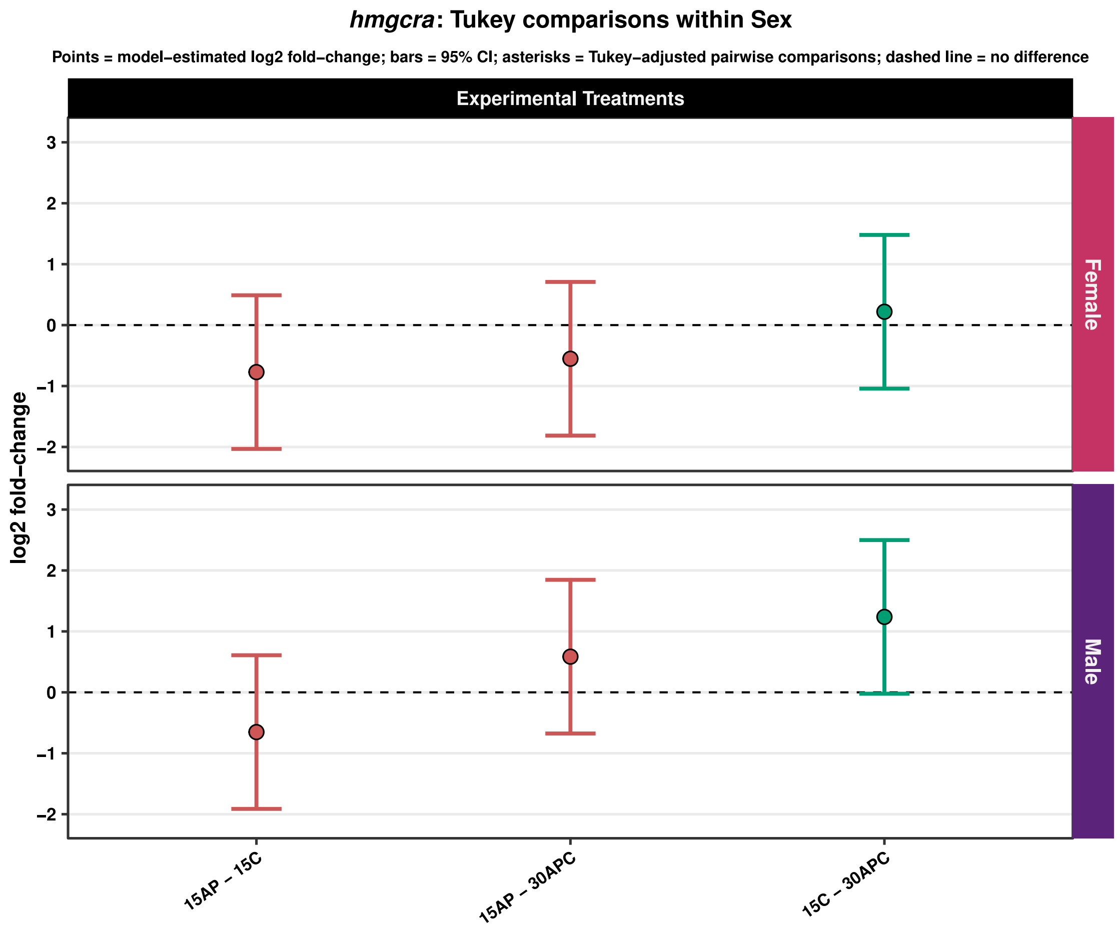


**Fig. S23:** Intestinal *hmgcra* gene expression across experimental diet comparisons and faceted by sex.

Diet pairwise contrasts of *hmgcra* expression. Each point is the model-estimated log2FC, and the vertical error bar is its 95% confidence interval.

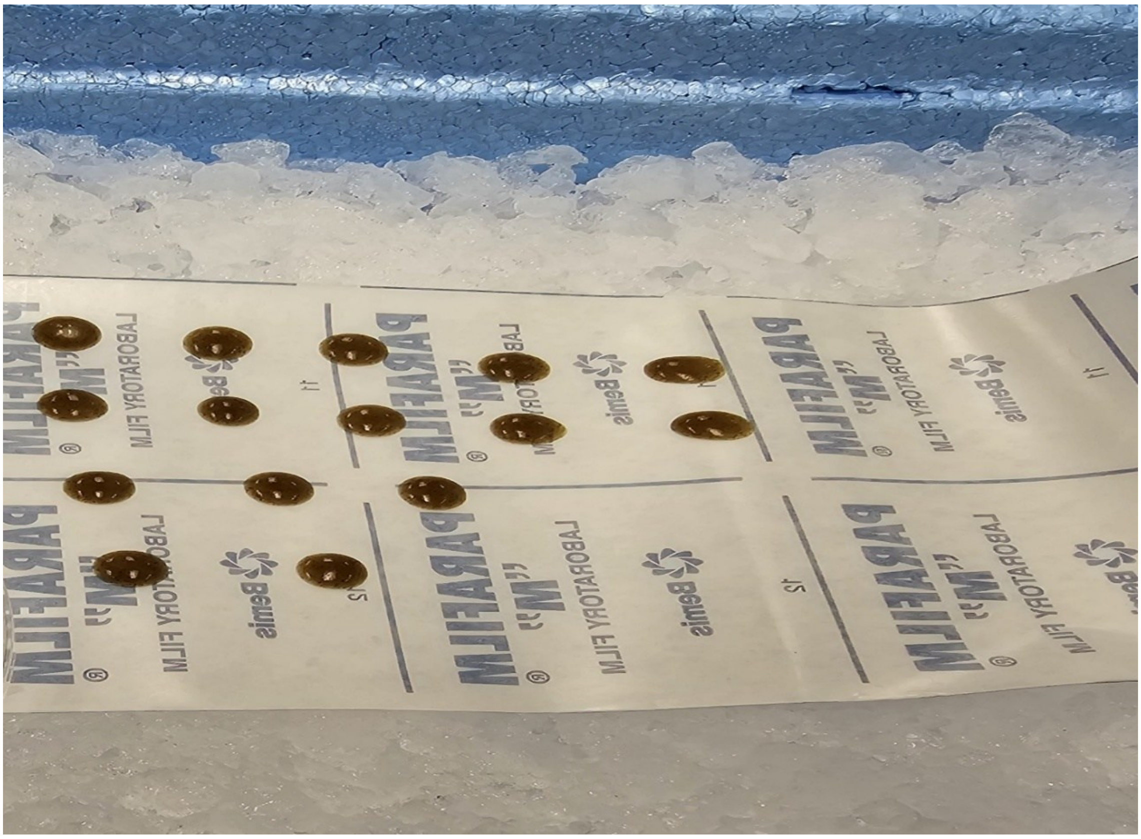

**Fig. S24:** Preparation of gel food.

The warm Repashy Community Plus gel (with or without saponin) was pipetted onto a Parafilm placed on ice. The gel droplets were then stored in the fridge (4-6 °C) until use. Adapted from[1].

| Diet | Sex | n |
|---|---|---|
| Control | F | 7 |
| Control | M | 7 |
| 15AP | F | 7 |
| 15AP | M | 7 |
| 15C | F | 6 |
| 15C | M | 7 |
| 30APC | F | 7 |
| 30APC | M | 7 |

**Fig. S25:** Liver metabolomics sample sizes per diet and sex.

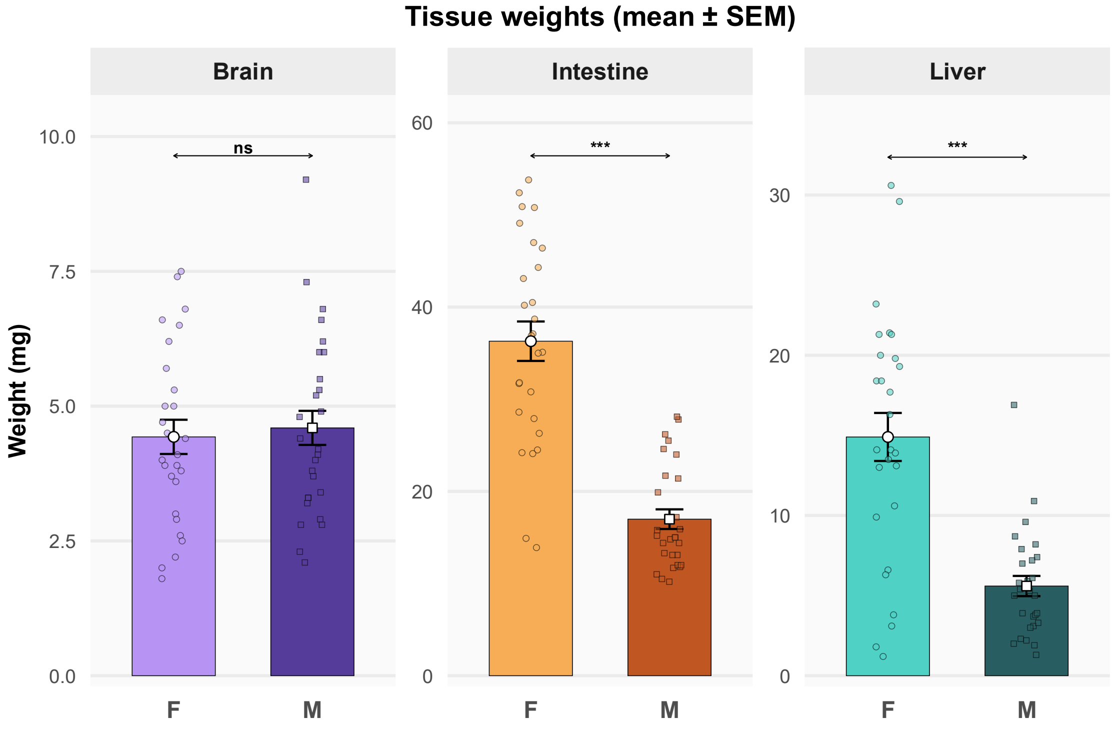


**Fig. S26:** Mean brain, intestine, and liver weights by sex.

Bar charts displaying the mean weights (mg) of the brain, intestinal, and liver samples in our study, grouped by sex. The centre dots and error bars indicate the mean ± standard error of the mean. Each individual dot represents a unique sample. F denotes females, and M denotes males. Welch's $t$-test: ns (non-significant) denotes $p > 0.05$ and ***denotes $p < 0.001$.

| Diet summary (mean ± SEM; tank = experimental unit) | | | | | | | |
|---|---|---|---|---|---|---|---|
| Diet | n | Initial mean weight (g) | Final mean weight (g) | Weight difference (g) | Percentage gain (%) | SGR (%/day) | Survival (%) |
| Control | 5 | 0.480 ± 0.023 | 0.656 ± 0.023 | 0.176 ± 0.038 | 38.37 ± 10.00 | 1.498 ± 0.338 | 100.00 ± 0.00 |
| 15AP | 5 | 0.372 ± 0.025 | 0.631 ± 0.032 | 0.258 ± 0.015 | 70.54 ± 6.21 | 2.530 ± 0.168 | 100.00 ± 0.00 |
| 15C | 5 | 0.462 ± 0.026 | 0.632 ± 0.035 | 0.170 ± 0.036 | 37.94 ± 8.28 | 1.494 ± 0.306 | 100.00 ± 0.00 |
| 30APC | 5 | 0.459 ± 0.030 | 0.655 ± 0.039 | 0.195 ± 0.036 | 43.91 ± 8.82 | 1.695 ± 0.310 | 100.00 ± 0.00 |

**Fig. S27:** Growth parameters and survival percentage.

The main effect of "Diet" was not statistically significant on the weight difference ($F_{3,16} = 1.55$, $p = 0.24$), nor on the SGR ($F_{3,16} = 2.92$, $p = 0.07$). However, "Diet" had a modest effect on % weight gain ($F_{3,16} = 3.36$, $p = 0.045$). However, post hoc comparisons between diets were not significant ($p_{adj} > 0.05$). Survival % was uniform across the diets.

**Table S1:** qPCR primer pairs used in this study

| Gene | Primer sequence (5′-3′) | Melting Temp (Tm °C) | Efficiency (%) | Amplicon length | Accession number | Role | Ref |
|---|---|---|---|---|---|---|---|
| *actb1* | F: ATGGATGAGGAAATCGCTG<br>R: ATGCCAACCATCACTCCCTG | F: 62.7<br>R: 67.2 | 114 | 108 bp | BC045846.1 | Reference gene | 2 |
| *eef1a1a* | F: ATCGACCGTCGTTCTGGCAAG<br>R: AACGGTGTGATTTGAGGGAA | F: 70.6<br>R: 64.3 | 104.5 | 250 bp | AY422992.1 | Reference gene | 3 |
| *rpl13a* | F: TGGTGAGGTGTGAGGGTATCAAC<br>R: AATTTGCGTGTGGGTTTCAGAC | F: 67.1<br>R: 67 | 99 | 269 bp | NM_212784.1 | Reference gene | 3 |
| *tnfa* | F: GCTGGATCTTCAAAGTCGGGTGTA<br>R: TGTGAGTCTCAGCACACTTCCATC | F: 69.1<br>R: 67.8 | 104 | 107 bp | AY427649.1 | Immune response | 2 |
| *il1b* | F: CTCAGCCTGTGTGTTTGGGA<br>R: GGGACATTTGACGGACTCG | F: 66.4<br>R: 65.6 | 108.6 | 176 bp | NM_212844.2 | Immune response | 4 |
| *il6* | F: AGACCGCTGCCTGTCTAAAA<br>R: TTTGATGTCGTTCACCAGGA | F: 63.8<br>R: 64.2 | 106.5 | 107 bp | NM_001261449.1 | Immune response | 2 |
| *cldn1* | F: CTGCTGTATCTGTGGGAGTGAA<br>R: TAATCAGGAGAACAGGCGAAG | F: 64.2<br>R: 63.2 | 107.3 | 133 bp | NM_131770.2 | Intestinal permeability | 5 |
| *oclna* | F: AGATGTGGAGGACTGGGTCA<br>R: ATTACGGACGGGCAGAATC | F: 64.7<br>R: 63.8 | 101.9 | 119 bp | NM_212832.2 | Intestinal permeability | 5 |
| *muc2.1* | F: CTGTGCGGCTAAAGGATAATC<br>R: CTCTGTGAGGCTGGGCAATC | F: 62.4<br>R: 67.2 | 104.1 | 108 bp | XM_073942803.1 | Mucus secretion | 6 |
| *mmp9* | F: TGATGTGCTTGGACCACGTAA<br>R: ACAGGAGCACCTTGCCTTTTC | F: 66.4<br>R: 66.6 | 97.9 | 94 bp | NM_213123.1 | Degradation and remodelling of the extracellular matrix | 7 |
| *hmgcra* | F: TGAGCAGGTGGAGATGGAGT<br>R: GGGGAGTTGGGGGTTTGTC | F: 65<br>R: 67.7 | 99.4 | 104 bp | NM_001079977.3 | Biosynthesis of cholesterol | 8 |

The qPCR primer pairs were obtained from the literature. The melting points were obtained from the supplier (Sigma). The primer efficiencies were obtained from a seven-point dilution standard curve. The accession numbers were obtained by BLAST (Basic Local Alignment Search Tool). The amplicon lengths were obtained from Sanger sequencing (Source Genomics LTD, Cambridge).

**Table S2:** Calculated approximate composition of diets.

<table>
<tr><th></th><th colspan="2">Control basal*</th><th>15DF</th><th>30DF</th></tr>
<tr><td>Composition</td><td colspan="2">Fish Meal, Vital wheat gluten, Fish protein hydrolysed, Crustacea meal, Pea protein, Gelatine, Algal oil (Schizochytrium sp.), Fish oil.</td><td>Fish Meal, Vital wheat gluten, Fish protein hydrolysed, Crustacea meal, Pea protein, Gelatine, Algal oil (Schizochytrium sp.), Fish oil, 15% DF[!].</td><td>Fish Meal, Vital wheat gluten, Fish protein hydrolysed, Crustacea meal, Pea protein, Gelatine, Algal oil (Schizochytrium sp.), Fish oil, 30% DF[!].</td></tr>
<tr><td>Diet</td><td>Final DF inclusion**</td><td>Protein g/100 g final diet</td><td>Fat g/100 g final diet</td><td>Total carbohydrate (including DF) g/100 g final diet</td></tr>
<tr><td>Control</td><td>0%</td><td>62.0</td><td>13.0</td><td>7.0</td></tr>
<tr><td>15AP</td><td>13%</td><td>53.9</td><td>11.3</td><td>16.9 – 17.8</td></tr>
<tr><td>15C</td><td>13%</td><td>53.9</td><td>11.3</td><td>18.5 – 19.1</td></tr>
<tr><td>30APC</td><td>23.1%</td><td>47.7</td><td>10</td><td>25.9 – 27.3</td></tr>
</table>

*Nutritional additives: Vitamins, pro-vitamins and chemically well-defined substances having a similar effect (per kg): vitamin D3 (3a671) 3000 IU; vitamin E (3a700) 1.2 g; vitamin C (3a311) 1.9 g; betaine (3a925) 10.8 g. Compounds of trace elements (mg/kg): iron (3b103) 9; iodine (3b201) 0.8; zinc (3b605) 11.3; copper (3b405) 13.5; manganese (Eb502) 14.4. The exact percentage composition of ingredients is not disclosed by Sparos.** 15 g DF + 100 g basal = 13.0% w/w final diet and 30 g DF (15 g $DF_1$ + 15 g $DF_2$) + 100 g basal = 23.1% w/w final diet

[!]DF: The dietary fibres used were purchased as analytical-grade reagents rather than food-grade ingredients. Approximate composition values were estimated from supplier specifications/literature and expressed as g/100 g ingredient as supplied. The carbohydrate fraction was assumed to represent predominantly dietary fibre polysaccharides.

Apple pectin (Sigma): ~ 0 g protein, ~ 0 g fat, ~ 83-90 g total carbohydrate

Cellulose (Sigma): ~ 0 g protein, ~ 0 g fat, and ~ 95-100 g total carbohydrate

- Final DF-reagent inclusion = (30/130 x 100 = 23.1%)
- Protein = (62/130 x 100 = 47.7) g/100 g
- Fat = (13/130 x 100 = 10.0) g/100 g
- Carbohydrate minimum = $(7 + 15\times0.83 + 15\times0.95)/130 \times100 = 25.9$ g/100 g
- Carbohydrate maximum = $(7 + 15\times0.90 + 15\times1.00)/130 \times100 = 27.3$ g/100 g
